# Drug Repurposing Through a Bioinformatics Pipeline Applied on Fibrotic Diseases

**DOI:** 10.1101/2020.05.19.103945

**Authors:** Evangelos Karatzas, Andrea Kakouri, George Kolios, Alex Delis, George M. Spyrou

**Affiliations:** Deprt. of Informatics and Tele/coms, University of Athens, 15703, Athens, Greece; Deprt. of Bioinformatics, The Cyprus Inst. of Neurology and Genetics, 2370, Nicosia, Cyprus; The Cyprus School of Molecular Medicine, 2370, Nicosia, Cyprus; Deprt. of Medicine, Democritus University of Thrace, 68100, Alexandroupolis, Greece

**Keywords:** Idiopathic Pulmonary Fibrosis, Drug Repurposing, Drug Re-ranking, Pathway Analysis, Disease Similarity

## Abstract

**Subject:** Fibrotic diseases cover a spectrum of systemic and organ-specific maladies that affect a large portion of the population, currently without cure. The shared characteristic these diseases feature is their uncontrollable fibrogenesis deemed responsible for the accumulated damage in the susceptible tissues. *Idiopathic Pulmonary Fibrosis* (*IPF*), an interstitial lung disease, is one of the most common and studied fibrotic diseases and still remains an active research target.

**Objective:** We highlight unique and common (i) genes, (ii) biological pathways and (iii) candidate repurposed drugs among nine fibrotic diseases. We bibliographically explore the resulting candidate substances for potential anti-fibrotic mode of action and focus on diseases that appear to be more similar to *IPF* so as to jointly examine potential treatments.

**Methodology:** We identify key genes for the 9 fibrotic diseases by analyzing transcriptomics datasets. We construct gene-to-gene networks for each disease and examine these networks to explore functional communities of biological pathways. We also use the most significant genes as input in Drug Repurposing (DR) tools and re-rank the resulting candidates according to their structural properties and functional relationship to each investigated disease.

**Results:** We identify 7 biological pathways involved in all 9 fibrotic diseases as well as pathways unique to some of these diseases. Based on our DR results, we suggest captopril and ibuprofen that both appear to slow the progression of fibrotic diseases according to existing bibliography. We also recommend nafcillin and memantine, which haven’t been studied against fibrosis yet, for further wet-lab experimentation. We also observe a group of cardiomyopathy-related pathways that are exclusively highlighted for *Oral Submucous Fibrosis* (*OSF*). We suggest digoxin to be tested against *OSF*, since we observe cardiomyopathy-related pathways implicated in *OSF* and there is bibliographic evidence that digoxin may potentially clear myocardial fibrosis. Finally, we establish that *IPF* shares several involved genes, biological pathways and candidate inhibiting-drugs with *Dupuytren’s Disease*, *IgG4-related Disease*, *SSc* and *Cystic Fibrosis*. We propose that treatments for these fibrotic diseases should be jointly pursued.

## Introduction

Fibrotic diseases constitute a group of incurable maladies that are recognized by a fibrotic phenotype affecting various organs and tissues. Pertinent mechanisms escape the homeostatic signals and due to over-repairing, cause tissue scarring. The plethora and complexity of the underlying perturbations are responsible for the existing lack of treatments. Even though fibrotic diseases target various biological structures, they do share several underlying mechanisms (Wernig, et al., 2017; Wynn, 2007).

One of the most important fibrotic mechanisms is the extracellular matrix (ECM) deposition which is known to drive both the initiation as well as the progression of fibrogenesis (Herrera, *et al.*, 2018). ECM is a three-dimensional network of extracellular macromolecules such as collagen, enzymes and glycoproteins, that provide structural and biochemical support to surrounding cells (Theocharis, *et al.*, 2016). The uncontrolled accumulation of ECM macromolecules is responsible for the replacement of normal with non-functional fibrotic tissue. The transforming growth factor-β (TGF-β) cytokine is a key regulator of ECM, since TGF-β signals, and particularly SMAD proteins which are TGF-β signal transducers, act as transcription factors that induce ECM’s expression in mesenchymal cells (Verrecchia and Mauviel, 2007). Other common mechanisms among fibrotic diseases include the (i) bone morphogenic protein (BMP) signaling, (ii) overexpression of connective tissue growth factor (CTGF), (iii) Wnt/β-catenin pathway and (iv) platelet-derived growth factor (PDGF) signaling (Rosenbloom, *et al.*, 2017). In particular, PDGF-A-/PDGFRα signaling loops stimulate fibroblasts to synthesize ECM and release pro-fibrotic mediators (Bonner, 2004).

Although *Idiopathic Pulmonary Fibrosis* (*IPF*) is one of the most common and better studied fibrotic diseases, it still remains a very active research target. *IPF* is an interstitial lung disease (*ILD*), primarily involving the tissue and space around the air sacs of the lungs. *IPF* is of unknown etiology and due to rapid fibrotic progression, leads to death in about 3-5 years. According to (Hutchinson, *et al.*, 2015), data from 21 countries present an incident rate of 3-9 cases per 100,000 per year for North America and Europe and lower rates for East Asia (1.2–3.8 per 100,000) and South America (0.4–1.2 per 100,000). Recent developments have led to updates in the guidelines for *IPF* diagnosis. In (Raghu, *et al.*, 2018), a multidisciplinary committee provided new guidelines for *IPF* diagnosis by combining evidence from high-resolution computed tomography (HRCT) and histopathological patterns of ‘usual interstitial pneumonia (*UIP*)’, ‘possible *UIP*’ and ‘indeterminate for *UIP*’. The recommendations strongly advise against serum biomarker (MMP7, SPD, CCL18, KL6) measurements as an approach to distinguish *IPF* from other *ILDs* because of the high false-positive and false-negative result rates.

In a previous work (Karatzas, *et al.*, 2019), we analyzed fibrotic molecular and phenotypic data from the Malacards and Human Phenotype Ontology databases, for 14 fibrotic diseases in an initial effort to group fibrotic diseases and identify similarities with *IPF*. The diseases that we observed being more similar to *IPF* were *Silicosis*, *Cystic Fibrosis* (*CF*), *Systemic Sclerosis* (*SSc*) and *IgG4-related Disease*. Nevertheless, drawing conclusions regarding the similarities of the underlying mechanisms and pathogenesis among diseases still remains an open research challenge.

In this paper, we study transcriptomics datasets regarding 9 of the 14 fibrotic diseases from (Karatzas, *et al.*, 2019) and present common and unique (i) genes, (ii) biological pathways and (iii) candidate repurposed drugs among these fibrotic diseases. We undertake this endeavor in an effort to group similar fibrotic diseases, as well as to better understand and potentially treat fibrosis. To attain our objective, we analyze gene expression datasets from microarray and next generation RNA-Sequencing (RNA-Seq) experiments to identify over- and under-expressed genes related to each of the 9 designated diseases. We then engage respective gene lists to drive random walks on a functionally-connected network of biological pathways through our pathway analysis methodology, called PathWalks (Karatzas, et al., 2020), to identify key fibrotic pathway communities. We also use the gene lists as input into computational signature-based Drug Repurposing (DR) tools (Lussier and Chen, 2011) to help designate potential therapeutic options against fibrosis. Subsequently, we re-rank the repurposed drugs based on their drugability and functional relation to each disease (i.e., gene targets) through our CoDReS tool (Karatzas, *et al.*, 2019). We further screen the drug candidates by prioritizing compounds that are structurally dissimilar to drugs that have already failed in clinical trials against fibrotic diseases. This is justifiable since drugs with similar structures tend to have similar mode of action (Campillos, *et al.*, 2008). Finally, we explore the existing bibliography for evidence that further supports the potential anti-fibrotic action of our proposed drugs. We present a flowchart of our proposed analysis pipeline in Figure 1. It is in this context that we seek to establish common and unique fibrosis-related genes, pathways, and drugs in an effort to group fibrotic diseases for potential common treatments. Moreover, we identify unique characteristics among fibrotic diseases of interest so as our results can be independently verified through wet lab experimentation.

**Figure 1:**
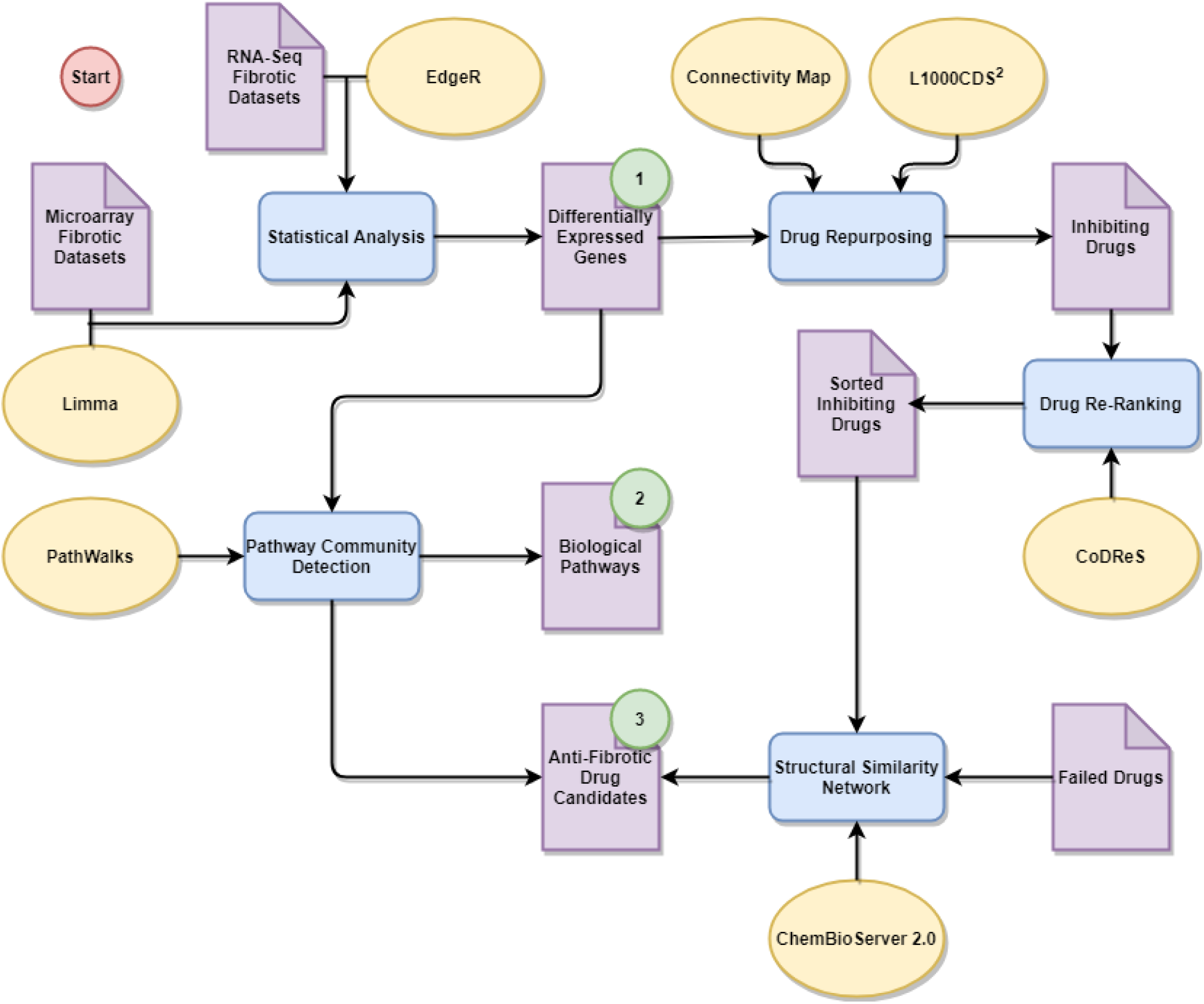
Flowchart of our pipeline regarding pathway analysis and drug repurposing for fibrotic diseases based on transcriptomics data. Purple pages represent data, blue rectangles represent procedures, yellow ellipses represent tools and methodologies and green circles indicate results regarding: (1) genes, (2) biological pathways and (3) inhibiting-drug candidates. We start by analyzing transcriptomics data from several tissues regarding 9 fibrotic diseases to extract key respective gene lists. We then identify common and unique perturbed biological pathways as well as potential therapeutic candidates by exploiting these gene lists.

## Materials and Methods

### Microarray Dataset Analysis

In the first step of the described pipeline we identify unique and common genes among fibrotic diseases by analyzing transcriptomics datasets. We then use these gene lists as input in pathway analysis and DR tools, to highlight biological pathways and candidate drugs regarding fibrosis. We initially utilize 14 microarray datasets regarding the 9 fibrotic diseases of Table 1 from Gene Expression Omnibus (GEO) (Edgar, *et al.*, 2002). We analyze the pre-processed *Series Matrix Files* of the data, provided by each of these 14 experiments. We apply the *normalizeQuantiles* function, from the Limma package (Ritchie, *et al.*, 2015) of the R programming language to non-normalized datasets. We then apply a log_2_ transformation to non-log_2_ datasets. If a dataset is already log_2_ transformed but not normalized, we remove the log_2_ transformation, then normalize and finally re-apply the log_2_ transformation. After executing the Limma statistical analysis, we use a significance cutoff-threshold (*p-value* < *0.05*) on the results. We remove any genes that have dual sign probes (over- and under-expressed at the same time). In the case of duplicate gene entries, we keep the result with the highest significance (i.e., lowest *p-value* score). Finally, we extract lists of the top-*150* over- and top-*150* under-expressed genes, based on fold change, per experiment which we then use as input in pathway analysis and DR tools.

**Table 1:** Dataset information regarding the GEO microarray experiments. We analyze 4 *IPF* microarray datasets where all samples are derived from lung tissue. *CF* samples come from 2 microarray datasets from rectal mucosal epithelia and nasal epithelial cells respectively. *SSc* samples are derived from skin tissue. The 2 *Myelofibrosis* datasets contain bone marrow and bone marrow stromal cell samples respectively. The *OSF* samples are derived from oral buccal mucosa. The *Dupuytren’s Disease* samples are derived from hand connective tissue. The *Polycystic Kidney Disease* samples are derived from renal cysts. The *IgG4-related Disease* samples come from labial salivary glands and finally, *Schistosomiasis* samples from liver tissue. Additionally, the number of samples (total, disease and control) that we experimented on is also provided for each dataset entry.

| Dataset ID | Disease | Tissue | #Samples | #Disease | #Control |
| --- | --- | --- | --- | --- | --- |
| GSE110147 | *IPF* | Lung | 33 | 22 | 11 |
| GSE53845 | *IPF* | Lung | 48 | 40 | 8 |
| GSE32537 | *IPF* | Lung | 207 | 167 | 50 |
| GSE35145 | *IPF* | Lung | 8 | 4 | 4 |
| GSE15568 | *CF* | Rectal mucosal epithelia | 29 | 16 | 13 |
| GSE40445 | *CF* | Nasal epithelial cells | 10 | 5 | 5 |
| GSE95065 | *SSc* | Skin | 33 | 18 | 15 |
| GSE124281 | *Myelofibrosis* | Bone marrow | 12 | 4 | 8 |
| GSE44426 | *Myelofibrosis* | Bone marrow mesenchymal stromal cells | 12 | 6 | 6 |
| GSE64216 | *OSF* | Oral buccal mucosa | 6 | 4 | 2 |
| GSE75152 | *Dupuytren’s* | Hand connective tissue | 24 | 12 | 12 |
| GSE7869 | *Polycystic Kidney* | Renal cysts | 21 | 18 | 3 |
| GSE40568 | *IgG4-related Disease* | Labial salivary gland | 8 | 5 | 3 |
| GSE61376 | *Schistosomiasis* | Liver | 10 | 6 | 4 |

We exclude any samples from the microarray datasets that do not correspond to the fibrotic diseases of this study. We provide some clarifications regarding these discarded samples: for the *Myelofibrosis* dataset GSE124281, we discard disease sample GSM3526859 since it was pooled from 2 patients and also exclude disease samples from patients with *Essential Thrombocythemia*. For the *Oral Submucous Fibrosis* (*OSF*) dataset GSE64216, we exclude the 2 *Squamous Cell Carcinoma* samples. We exclude 5 *Sjogren’s Syndrome* samples from the *IgG4-related Disease* dataset GSE40568. We also exclude 7 *Hepatitis B* samples from the *Schistosomiasis* dataset GSE61376. Finally, we exclude the *Non-Specific Interstitial Pneumonia* (*NSIP*) and *IPF-NSIP* samples from the *IPF* dataset GSE110147. We also note that in the *CF* dataset GSE15568 patients carry the CF-specific D508 mutated CFTR-allele while in the *CF* dataset GSE40445 patients have a confirmed F508del homozygosity for the CFTR gene. Table 1 shows dataset details regarding diseases, sample tissues and number of samples (total, disease and control) after our exclusions.

### RNA-Seq Dataset Analysis

Complementary to the microarray dataset analysis, we use RNA-Seq datasets from GEO that are available for *IPF*, *CF* and *SSc*. We analyze the pre-processed *Raw Count Matrix Files* of the data provided by each experiment using R’s EdgeR package (Robinson, *et al.*, 2010). We perform normalization on the gene count matrices for RNA composition, by calculating the normalizing factors for the library sizes using the *trimmed mean of M-values* method between each pair of samples. We keep the genes with a minimum requirement of *1 count per million (CPM)* across 2 or more libraries for each group (affected and control). Once we normalize the data, we then test for differential expression between patient and healthy control samples using the *quasi-likelihood F-test (QLF)*; this is a statistical method based on negative binomial distribution that calculates the natural variation between biological replicates (Chen, *et al.*, 2016). We then use a significance cutoff-threshold on the results (*p-value = 0.05*). In agreement with the microarray dataset analysis procedure, we extract lists of the top-*150* over- and top-*150* under-expressed genes per experiment, based on fold change.

Again, we exclude samples that do not correspond to the fibrotic diseases of this study. Regarding the GSE124548 *CF* dataset, we exclude the post-drug patient samples and compare pre-drug patient and healthy control samples. Similarly, for the *IPF* dataset GSE99621 we exclude the macroscopically unaffected (normal-appearing) patient samples and test for differentially expressed genes between macroscopically affected and healthy controls. Table 2 shows dataset details regarding diseases, sample tissues and number of samples (total, disease and control) after the exclusions.

**Table 2:** Dataset information regarding the GEO RNA-Seq experiments. We analyze 2 RNA-Seq datasets containing *IPF* lung samples. We also analyze 1 RNA-Seq dataset from *CF* whole blood samples and 1 RNA-Seq dataset of *SSc* samples from monocyte-derived macrophages. Additionally, the number of samples (total, disease and control) that we experimented on, for each dataset entry, is shown on the table.

| Dataset ID | Disease | Tissue | #Samples | #Disease | #Control |
| --- | --- | --- | --- | --- | --- |
| GSE99621 | IPF | Lung | 16 | 8 | 8 |
| GSE134692 | IPF | Lung | 72 | 46 | 26 |
| GSE124548 | CF | Whole blood | 40 | 20 | 20 |
| GSE104174 | SSc | Monocyte-derived macrophages | 72 | 15 | 57 |

### Pathway Community Detection

We aggregate the key differentially expressed genes (over- and under-expressed) among experiments per disease. We utilize these gene lists in conjunction with our PathWalks methodology (Karatzas, *et al.*, 2020) to detect biological pathways and their functional communities that are unique or common among the 9 fibrotic diseases of this study. PathWalks is a pathway analysis and community detection methodology that guides a walker on a pathway-to-pathway network of functional connections exploiting an integrated network of genetic information that we make available.

We construct 9 gene-to-gene networks (1 per fibrotic disease) based on the extracted gene lists from the microarray and RNA-Seq datasets analysis procedure. We achieve this by using these gene lists as input in Cytoscape’s (Shannon, *et al.*, 2003) GeneMANIA plug-in (Montojo, *et al.*, 2010). Specifically, we execute GeneMANIA with the following options: (i) *0* resultant genes and attributes (i.e., not adding enriched genes or functional categories), (ii) weight assigned based on query gene and (iii) network types based on the default selections for gene co-expression, physical interaction and pathways. We export the “*normalized max weight*” column of GeneMANIA’s results that contains the relation weights between gene pairs. We then aggregate these weight-values among the different types of networks for the same gene pair to emphasize the relevance of the connection between the gene pair, and finally, we translate the gene symbols to their respective identifier from the KEGG database (Kanehisa, *et al.*, 2017) to match them with the respective biological pathways they participate in.

In more detail, the GeneMANIA results present a weight for each participating type of network and a separate weight for each edge. GeneMANIA’s execution algorithm consists of two parts; a label propagation function which assigns weights on the edges of the composite network and a ridge regression (regularized linear regression) function which assigns weights to the various types of networks (Mostafavi, *et al.*, 2008). The final composite network is a weighted sum of the individual data sources. The participating networks’ weights sum to *100*% and reflect the relevance of each data source for predicting membership in the query list (Warde-Farley, *et al.*, 2010). Edges are weighted relatively to the co-functionality between genes stated in each dataset. The final edge values are weighted sums of the respective normalized weights multiplied by the respective network’s weight (Zuberi, *et al*., 2013). We note that each entry of GeneMANIA’s “*normalized max weight*” column describes the normalized value for the maximum raw edge weight of a gene pair, among its various edge-entries for the same network (i.e., multiple scores from different databases for the same gene pair and network type).

Apart from the gene-to-gene networks that we implement for the PathWalks execution, a pathway-to-pathway network is also required by the algorithm. We create a network of biological pathways based on their functional relations according to the KEGG database. We assign edge scores equal to the number of common genes between each pair of pathways. We exploit the pathway-to-pathway network in all 9 executions (1 per disease) and finally, compare the pathway results of each disease associated to the respective genetic information networks.

The pathway nodes and their functional communities that are highlighted by PathWalks tend to favor hubs with high betweenness centrality and strength scores, as our algorithm is based on shortest paths while traversing the network of pathways. The betweenness centrality metric represents a node’s participation in the total shortest paths of a network while the strength metric represents a node’s sum of edge weights. We are particularly attentive to the most traversed pathways that are not necessarily highlighted due to topology but mainly through the guidance of the genetic information map. We conclude our pathway analysis by identifying pathways that are either common or uniquely perturbed in fibrotic diseases.

### Drug Repurposing

We use the differentially expressed gene lists from each of the 14 microarray and 4 RNA-Seq experiments as input in 2 signature-based DR tools: (1) Connectivity Map (CMap) (Subramanian, *et al.*, 2017) and (2) L1000CDS^2^ (Duan, *et al.*, 2016). We utilize the top-*150* over- and top-*150* under-expressed genes from each experiment, as CMap suggests each input gene list is between *10* and *150* genes for optimal results. We need the resultant substances to inverse the input genes’ (disease) signature. To achieve this in CMap, we choose the detailed list results, then select the “*Compound*” choice of the “*Perturbagen type*” subset and sort the results by ascending connectivity score. In L1000CDS^2^ we select the “*reverse*” option in the “*Configuration*” settings on the query page since, again, we require drugs that reverse the input disease signature. We keep substances with summary-inhibition score less than −*50* (−*100* denotes maximum inhibition) from each CMap experiment and all 50 returned entries from L1000CDS^2^.

We then use our drug re-ranking tool CoDReS (Karatzas, et al., 2019) to screen the lists of drug candidates and identify the most promising inhibitors for each fibrotic disease. The tool integrates information from biological databases to calculate functional associations among input drugs and queried diseases along with their binding affinities. CoDReS also calculates the potential drugability of each drug according to the Lipinski and Veber rules (Lipinski, et al., 1997; Veber, et al., 2002). This tool constructs structural clusters of the input substances, based on their chemical composition, by utilizing the affinity propagation algorithm (Bodenhofer, *et al.*, 2011) and proposes the top-ranked inhibitors per cluster for further testing against a disease. We compile an aggregated list of repurposed candidate drugs, derived from both DR tools across all experiments, for each disease and use them as input in CoDReS while selecting the respective disease for each execution.

To further screen the results, we calculate the structural similarities among the short drug lists from CoDReS and substances that have previously failed in clinical trials against fibrotic diseases. To achieve this, we construct a structural similarity network of our re-ranked drugs and the failed drugs of fibrotic diseases as found in repoDB (Brown and Patel, 2017) (last update: July 28, 2017). We use an edge cutoff-threshold of *substance similarity = 0.7*. With this approach, we avoid prioritizing candidates with a lower chance of success for further experimentation, since drugs with similar chemical structures tend to have similar mode of action. We construct the structural similarity network through the ChemBioServer 2.0 web tool (Karatzas, et al., 2020) and visualize it with Cytoscape (Shannon, *et al.*, 2003).

Subsequently, we designate the most promising anti-fibrotic candidates while focusing on *IPF*, by identifying the gene targets of the re-ranked drugs that participate in key implicated biological pathways. We utilize drug-target information from the DrugBank (Wishart, *et al.*, 2018), DrugCentral (Ursu, *et al.*, 2019) and DGIdb (Cotto, *et al.*, 2018) databases and pathway-gene information from the KEGG pathway database (Kanehisa, *et al.*, 2017). We finally explore the existing bibliography for anti-fibrotic indications concerning our proposed drug candidates to suggest the most promising for further *in vitro* and *in vivo* testing.

## Results

### Key Fibrosis-related Genes

In this section, we present key fibrotic genes we identify through our microarray and RNA-Seq analyses on pertinent datasets. We aggregate the top-150 over- and top-150 under-expressed gene lists among same disease experiments and observe that *IPF* has the most common over-expressed genes with *Dupuytren’s Disease* (*35*) and the most common under-expressed genes with *Myelofibrosis* (*28*). *Schistosomiasis* has the least number of common gene entries with *IPF* (*3* over- and *3* under-expressed). We identify the *LCN2* gene being over-expressed in *IPF*, *CF*, *Schistosomiasis* and *SSc* and the *FBLN1* gene being under-expressed in *CF*, *Myelofibrosis*, *Polycystic Kidney Disease* and *SSc* and over-expressed in *IgG4-related Disease*.

*LCN2* encodes for the lipocalin-2 protein which is associated with neutrophil gelatinase (Kjeldsen, *et al.*, 1993) and is known to limit bacterial growth (Flo, et al., 2004). Takahashi *et al.* study the *LCN2* expression in (i) the skin of patients with *SSc*, (ii) bleomycin-treated mice, and (iii) Fli1-deficient endothelial cells. Their experiments show that *LCN2* is associated with dermal fibrosis in early *Diffuse Cutaneous SSc* cases (some of which are also diagnosed with *ILD*) but not with *ILD* markers such as diffusion lung capacity for carbon monoxide or vital capacity (Takahashi, *et al.*, 2015). Their results show correlation between *LCN2* and progressive skin sclerosis as well as pulmonary vascular involvement that leads to pulmonary arterial hypertension in *SSc*. Furthermore, (Nakagawa, *et al.*, 2015) demonstrate increased expression of *LCN2*, correlated with tubulointerstitial fibrosis and tubular cell injury in patients with *Chronic Kidney Disease*. In another study, Kim *et al.* show increased *LCN2* levels in urine samples of patients with *Chronic Hepatitis C* accompanied by hepatic fibrosis (Kim, et al., 2010).

Other studies show no direct correlation between *LCN2* and hepatic fibrosis. Regarding *Nonalcoholic Fatty Liver Disease* (*NAFLD*), Milner *et al.* observe no association between *LCN2* and steatosis, lobular inflammation, ballooning or fibrosis even though the expression of *LCN2* is significantly elevated in *NAFLD* samples compared to controls (Milner, *et al.*, 2009). Furthermore, Borkham-Kamphorst *et al.* study the expression levels of *LCN2* in rat models with acute and chronic liver injury. Their results show correlation of *LCN2* to liver damage and resulting inflammatory responses but not to the degree of liver fibrosis (Borkham-Kamphorst, *et al.*, 2011).

Through our analysis, we observe that *LCN2* is over-expressed in various fibrotic diseases and their respective targeted tissues, such as *SSc* and the affected skin tissue. However, the correlation between the over-expression of *LCN2* and fibrosis is not yet clear and its role might differ among the various fibrotic diseases and affected tissues.

The *FBLN1* gene encodes for the fibulin-1 protein which is an ECM and plasma glycoprotein (Argraves, *et al.*, 1990). Chen *et al.* study the effects of TGF-β1 on the regulation of *FBLN1* on primary human airway smooth muscle cells from volunteers with and without *Chronic Obstructive Pulmonary Disease* (*COPD*) where small airway fibrosis occurs. They show that TGF-β1 causes a decrease in *FBLN1* mRNA and soluble FBLN1 while it increases the FBLN1 in ECM due to the sequestration of soluble FBLN1 (Chen, *et al.*, 2013). This fact can partially justify the under-expression of *FBLN1* in 4 diseases of our transcriptomics analyses (even though it was over-expressed in *IgG4-related Disease*), since gene expression is measured by the level of the corresponding mRNA present in a cell.

(Liu, *et al.*, 2016) suggest that *FBLN1* may be involved in the pathogenesis of not only *COPD* but also of asthma and *IPF*. Specifically, (Liu, *et al.*, 2016) show that the deletion of *FBLN1*’s variant *FBLN1C* in mouse models inhibits airway and lung remodeling in chronic asthma and lung fibrosis while also protects against *COPD*. The *FBLN1C* variant is known to increase the proliferation of lung fibroblasts in *COPD* and *IPF* patients (Ge, *et al.*, 2015). Regarding *IPF*, (Jaffar, *et al.*, 2012) identify increased soluble FBLN1 levels in the serum of patients even though there is no overexpression of *FBLN1*’s deposition in the airway tissue. Finally, Hansen *et al.* study *FBLN1* with regard to myocardial fibrosis. In (Hansen, *et al.*, 2013), they suggest that, since *Aortic Valve Stenosis* (*AVS*) causes cardiac fibrosis and since they observe elevated expression of the *FBLN1* gene in *AVS* samples against controls, the *FBLN1* must be expressed as part of the fibrotic process. Even though *FBLN1* is undeniably implicated in various fibrotic diseases, its expression levels in different tissues and its actual role regarding different fibrotic diseases should be further tested.

We show the 2 common-gene participation networks (over/under-expressed) in Figure 2. Table 3 depicts the top-3 over- and top-3 under-expressed significant (p-value < .05) genes of each experiment based on their log fold change (logFC) values. Supplementary Tables 1 and 2 list all top common and unique genes (over- and under-expressed respectively), regarding the 9 fibrotic diseases.

**Figure 2:**
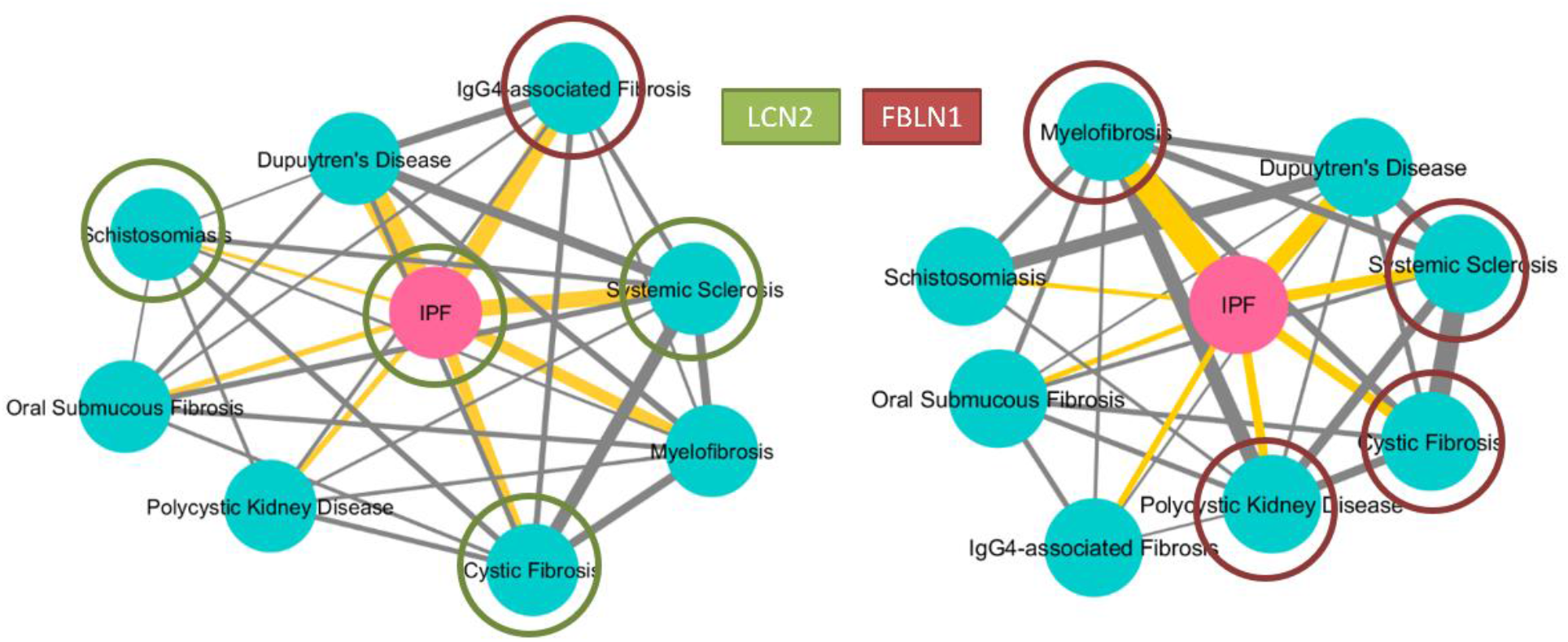
Common-gene participation networks. On the left network, the edge values are relative to the number of common over-expressed genes between pairs of fibrotic diseases. *IPF* has *35* common over-expressed genes with *Dupuytren’s Disease* followed by *23* common with *IgG4-related Disease* and *20* with *SSc*. Our analyses indicate that the *LCN2* gene is over-expressed in *IPF*, *CF*, *Schistosomiasis* and *SSc*. On the right network, the edge values are relative to the number of common under-expressed genes between pairs of fibrotic diseases. *IPF* has *28* common under-expressed genes with *Myelofibrosis* followed by *16* with *Dupuytren’s Disease* and *10* with *SSc.* Our analyses indicate that the *FBLN1* gene is under-expressed in *CF*, *Myelofibrosis*, *Polycystic Kidney Disease* and *SSc* but over-expressed in *IgG4-related Disease*. *Schistosomiasis* has only *3* common over- and *3* common under-expressed genes with *IPF*.

**Table 3:** Top-3 over- and top-3 under-expressed genes of each experiment based on their logFC values. We acquire gene symbols from GEO, from the respective platform-translation files of each experiment.

| Dataset ID | Disease Name | Experiment Type | Upregulated Gene | logFC | Downregulated Gene | logFC |
| --- | --- | --- | --- | --- | --- | --- |
| GSE110147 | IPF | Microarray | *MMP1* | 4.08 | *VTRNA1-1* | -5.63 |
|  |  |  | *SPP1* | 3.96 | *RNA5SP242* | -5.22 |
|  |  |  | *BPIFB1* | 3.85 | *SNORD41* | -4.33 |
| GSE53845 | IPF | Microarray | *MMP1* | 4.68 | *PNMT* | -3.97 |
|  |  |  | *UBD* | 4.22 | *IL1RL1* | -3.40 |
|  |  |  | *MMP7* | 4.08 | *DEFA3* | -3.26 |
| GSE32537 | IPF | Microarray | *BPIFB1* | 2.99 | *IL1R2* | -2.81 |
|  |  |  | *ASPN* | 2.63 | *MGAM* | -2.17 |
|  |  |  | *MMP7* | 2.56 | *FCN3* | -2.14 |
| GSE35145 | IPF | Microarray | *SFRP2* | 3.67 | *IL1R2* | -4.13 |
|  |  |  | *MMP7* | 3.55 | *S100A12* | -3.19 |
|  |  |  | *LOC649923* | 3.48 | *S100A8* | -3.13 |
| GSE99621 | IPF | RNA-Seq | *TTR* | 5.77 | *CHIAP2* | -4.90 |
|  |  |  | *OR51E1* | 5.75 | *WFDC12* | -4.72 |
|  |  |  | *GTF2H4* | 5.73 | *PNMT* | -4.67 |
| GSE134692 | IPF | RNA-Seq | *ADIPOQ* | 10.00 | *DEFA1B* | -8.99 |
|  |  |  | GP1BB | 7.89 | OLFM4 | -7.50 |
|  |  |  | CST1 | 6.83 | DEFA4 | -6.95 |
| GSE15568 | CF | Microarray | ALDOB | 3.08 | BRINP3 | -2.29 |
|  |  |  | ISG20 | 2.39 | SNRK | -1.70 |
|  |  |  | TNFRSF11B | 2.10 | TLR7 | -1.67 |
| GSE40445 | CF | Microarray | MALAT1 | 2.60 | PROS1 | -2.39 |
|  |  |  | LY6D | 2.59 | CRISP2 | -2.33 |
|  |  |  | CSF2RB | 2.25 | FABP6 | -2.32 |
| GSE124548 | CF | RNA-Seq | LOC107987345 | 4.46 | RPL10P6 | -2.76 |
|  |  |  | BCORP1 | 4.45 | RPL10P9 | -1.90 |
|  |  |  | TTTY15 | 3.91 | LOC102723846 | -1.80 |
| GSE95065 | SSc | Microarray | COL4A4 | 4.68 | ZBTB7C | -3.53 |
|  |  |  | SFRP4 | 3.16 | GSN | -3.22 |
|  |  |  | CXCL13 | 2.97 | SGCA | -3.12 |
| GSE104174 | SSc | RNA-Seq | MTND1P23 | 6.32 | LINC00221 | -3.65 |
|  |  |  | HBA2 | 6.10 | DOK5 | -2.74 |
|  |  |  | CRISP3 | 4.18 | KCNJ10 | -2.16 |
| GSE124281 | Myelofibrosis | Microarray | IFI27 | 3.12 | LOC389342 | -1.72 |
|  |  |  | RAP1GAP | 2.29 | LOC286444 | -1.71 |
|  |  |  | IL1B | 2.03 | VCAN | -1.66 |
| GSE44426 | Myelofibrosis | Microarray | OXTR | 4.33 | HAND2 | -7.13 |
|  |  |  | PTPRB | 4.15 | CD70 | -6.08 |
|  |  |  | TRPC4 | 3.95 | CHI3L2 | -5.96 |
| GSE64216 | OSF | Microarray | MB | 10.12 | KRT76 | -6.73 |
|  |  |  | MYLPF | 10.03 | LOC649923 | -6.24 |
|  |  |  | CKM | 9.81 | IGLL3 | -6.07 |
| GSE75152 | Dupuytren's | Microarray | THBS4 | 5.94 | LEP | -4.65 |
|  |  |  | LOC649366 | 4.97 | PLIN | -4.39 |
|  |  |  | TNC | 4.68 | PPP1R1A | -4.36 |
| GSE7869 | Polycystic Kidney | Microarray | UMOD | 6.84 | SFRP2 | -5.97 |
|  |  |  | SLC7A13 | 6.43 | CTHRC1 | -5.62 |
|  |  |  | MIOX | 6.41 | SFRP4 | -5.49 |
| GSE40568 | IgG4-related Disease | Microarray | ADH1B | 3.93 | FOS | -5.56 |
|  |  |  | SPP1 | 3.73 | IGHM | -3.68 |
|  |  |  | HLA-DRB4 | 3.23 | IFI6 | -3.57 |
| GSE61376 | Schistosomiasis | Microarray | FAM100A | 2.85 | RPS4Y1 | -3.39 |
|  |  |  | IGFBP2 | 2.58 | GSTM2 | -3.33 |
|  |  |  | PTMS | 2.01 | GSTM1 | -2.89 |

### Identifying Fibrotic Pathway Communities

We use the differentially expressed gene lists per disease, to drive our pathway analysis methodology. The PathWalks algorithm favors pathway-nodes with high betweenness centrality and strength scores. PathWalks, which is based on random walks and utilizes shortest paths, highlights such nodes. In Figure 3 we depict biological pathways which are more likely to appear in the PathWalks results due to the topology of the network rather than pathways with direct biological association to each fibrotic disease. Specifically, the depicted orange nodes belong to the top-*5*% of pathways with the highest betweenness centrality. Pink nodes belong to the top-*5*% of pathways with the highest degree. Red nodes belong to both groups and yellow nodes depict pathways that do not belong to any of the above 2 groups. Yellow nodes still reach the top-*5*% of the PathWalks results with random pathway selection, due to their high functional connectivity edge values. Since we construct the pathway-to-pathway network based on functional connections, all colored pathways may be sensitive to biological perturbations in most use cases.

**Figure 3:**
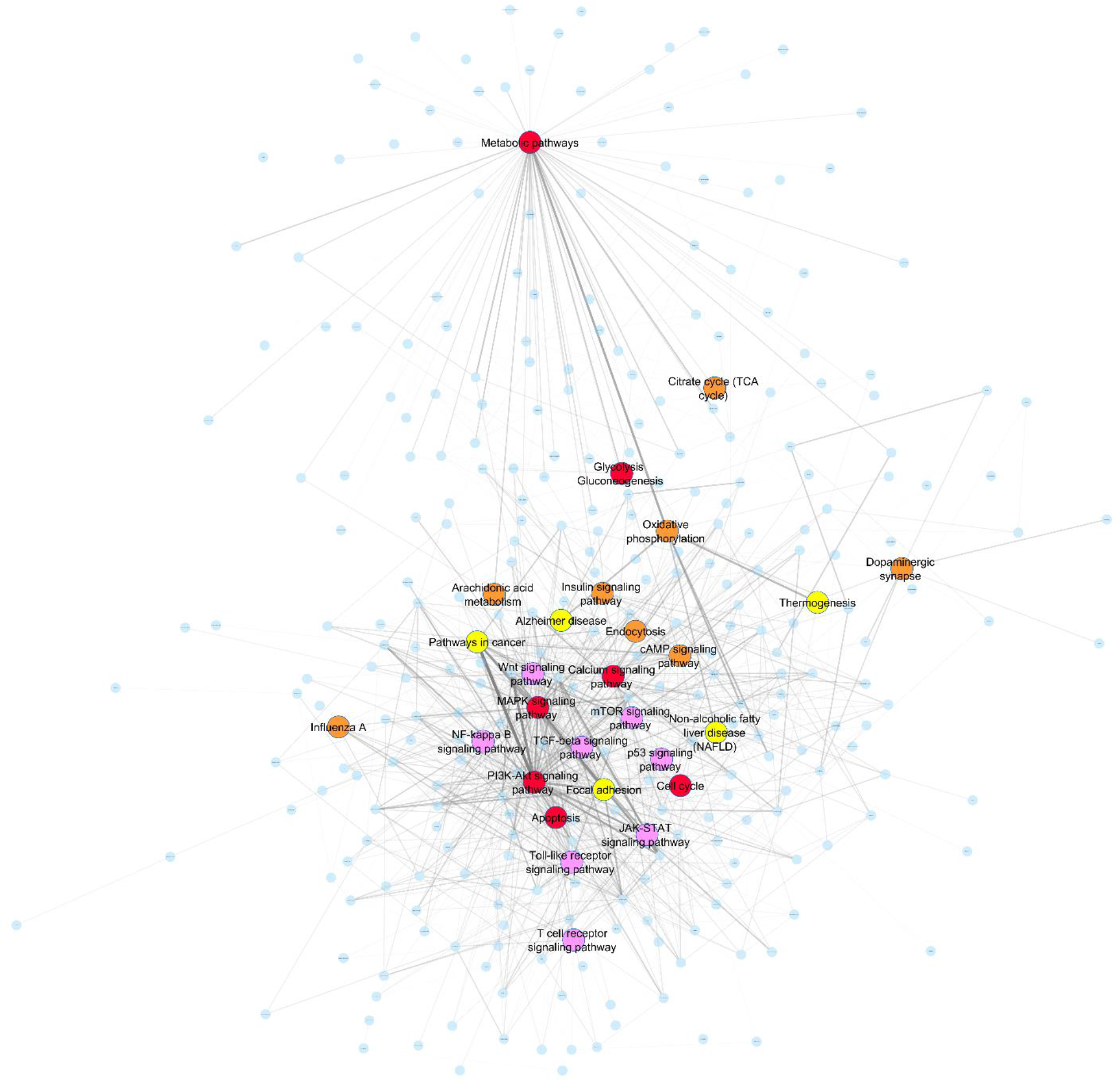
The base pathway-to-pathway network component that we use in the PathWalks execution. An edge indicates a functional connection between two pathways according to KEGG, and its thickness is relative to the number of their common genes. Orange nodes belong to the top-*5*% of pathways with the highest betweenness centrality, pink nodes to the top-*5*% with the highest degree and red nodes to both groups. Yellow nodes depict pathways that do not belong in any of these groups but still reach the top-*5* % in the results of a no-map PathWalks execution (i.e., random pathway selections) due to their high functional connectivity edge values. All colored nodes are more likely to appear in the results due to their topological characteristics rather than due to direct relation with the genetic information networks of each disease.

We identify disease-specific as well as common implicated pathways among the 9 fibrotic maladies through the PathWalks executions. In Figure 4 and in green color, we highlight nodes that reach the top-*5*% of the PathWalks results for at least 1 disease and do not belong in the topology-favored groups of Figure 3. We observe that not every topologically-favored node is highlighted by PathWalks. Therefore, we believe that even the topologically-favored nodes that are highlighted by the algorithm are not necessarily false-positive entries and should be considered for further experimentation regarding fibrosis. We then focus on the PathWalks-highlighted pathways of all 9 fibrotic diseases and draw the respective convex hulls per disease in Figure 5. We observe *7 common pathways across the 9 diseases*, all of which are favored by the topology. The pathways in question are “Metabolic”, “Cancer”, “MAPK signaling”, “PI3K-Akt signaling”, “Non-alcoholic fatty liver disease”, “Oxidative phosphorylation” and “Calcium signaling” pathways.

**Figure 4:**
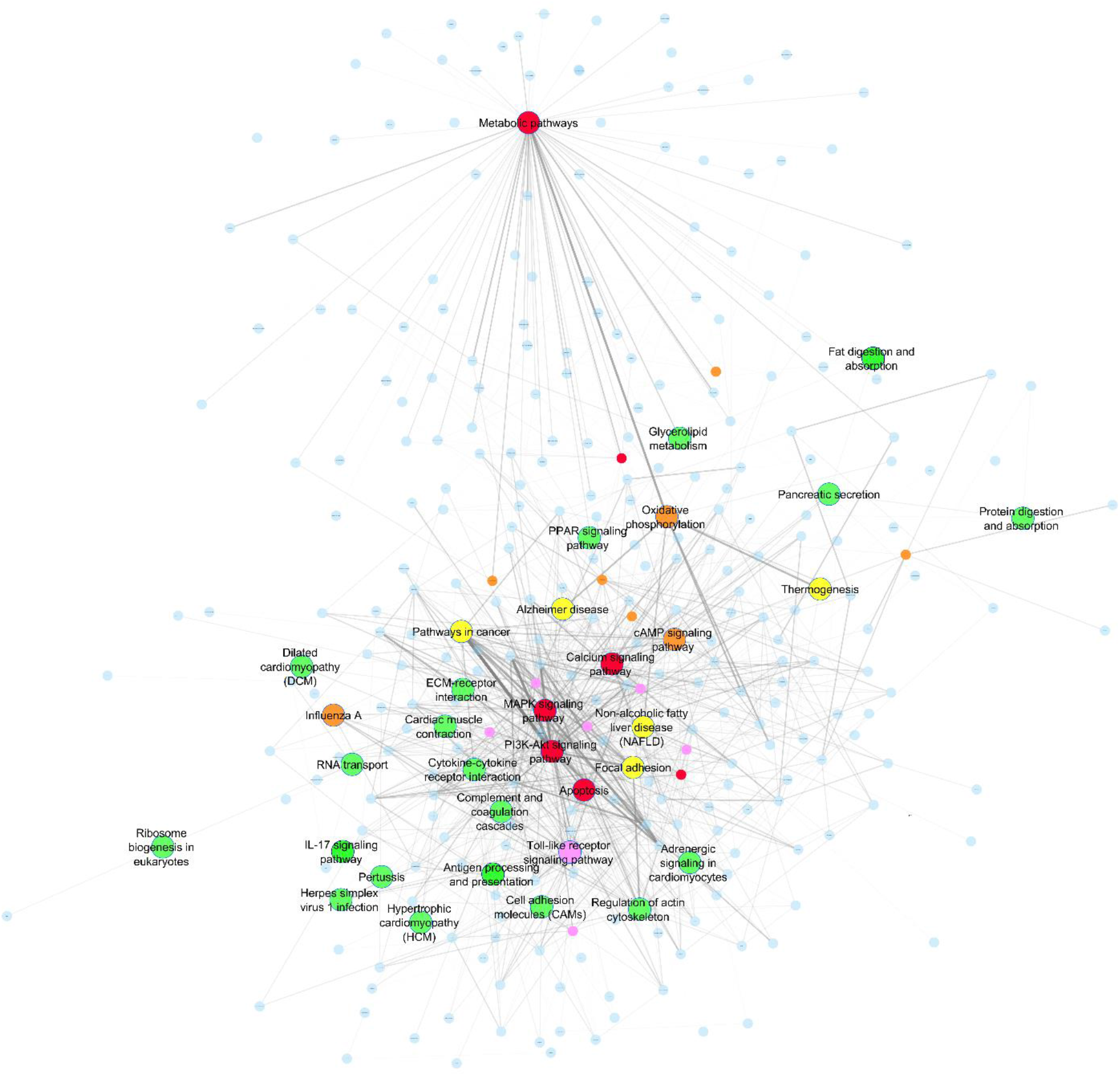
PathWalks results annotated on the basis of Figure 3 pathway-to-pathway. We highlight nodes that are exclusively based on the guidance of the gene maps in green color. We label all nodes that were produced in the top-*5*% of any PathWalks experiment. We observe that not every topologically-favored node is highlighted by PathWalks. Hence, we believe that even the topologically-favored nodes that are highlighted by the algorithm are not necessarily false-positive entries and should be further explored regarding fibrosis.

**Figure 5:**
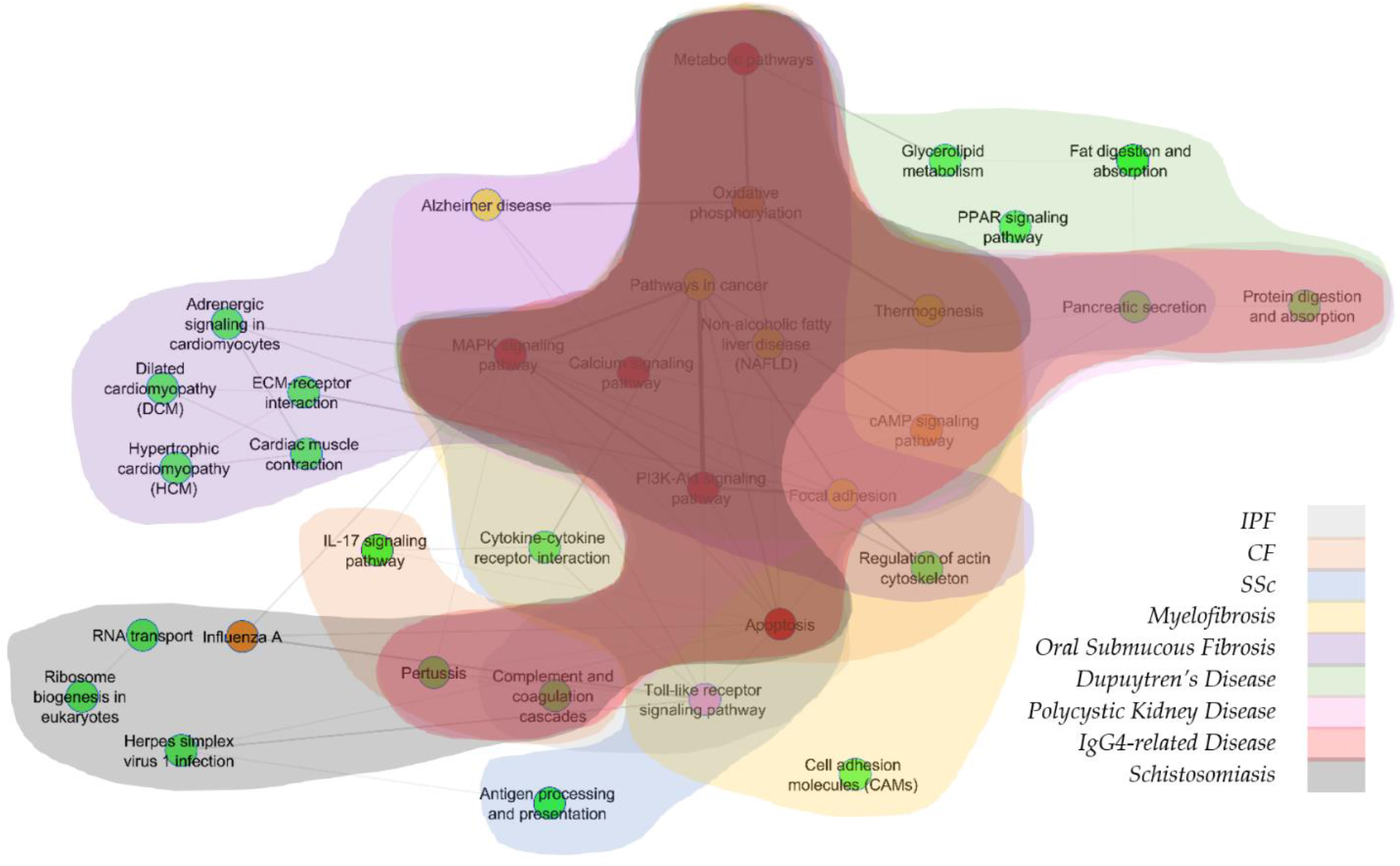
Convex hulls for the 9 fibrotic diseases highlighted pathways as produced by PathWalks. We observe that 7 pathways are common for all diseases including “Metabolic pathways”, “Pathways in cancer”, “MAPK signaling pathway”, “PI3K-Akt signaling pathway”, “Non-alcoholic fatty liver disease”, “Oxidative phosphorylation” and “Calcium signaling pathway”. We also identify unique highlighted pathways for some of the diseases. The “Cell adhesion molecules” pathway which belongs to the yellow convex hull is exclusive for *Myelofibrosis*. The “Antigen processing and presentation” pathway is included in the blue convex hull and is unique for *SSc*. The “IL-17 signaling” pathway is included in the orange convex hull and is unique for *CF*. “Fat digestion and absorption”, “PPAR signaling” and “Glycerolipid metabolism” pathways are unique for *Dupuytren’s Disease* and belong to the green convex hull. “Influenza A”, “RNA transport”, “Ribosome biogenesis in eukaryotes” and “Herpes simplex virus 1 infection” pathways belong to the dark gray convex hull and are unique for *Schistosomiasis*. Finally, the purple convex hull contains the “Dilated cardiomyopathy”, “Cardiac muscle contraction”, “Hypertrophic cardiomyopathy”, “Adrenergic signaling in cardiomyocytes” and “ECM-receptor interaction” pathways which are unique for *OSF*. We observe that 4 out of the 5 unique pathways of *OSF* are related to cardiomyopathies hinting for a novel approach of potential treatments since a link between OSF and cardiomyopathies is missing from the bibliography.

We also observe unique PathWalks-highlighted pathways in some of the fibrotic diseases. The “Cell adhesion molecules” pathway is exclusive for *Myelofibrosis*, the “Antigen processing and presentation” is exclusive for *SSc* and the “IL-17 signaling” is unique for *CF*. “Fat digestion and absorption”, “PPAR signaling” and “Glycerolipid metabolism” pathways are unique for *Dupuytren’s Disease*. “Influenza A”, “RNA transport”, “Ribosome biogenesis in eukaryotes” and “Herpes simplex virus 1 infection” pathways are unique for *Schistosomiasis*. Finally, the “Dilated cardiomyopathy”, “Cardiac muscle contraction”, “Hypertrophic cardiomyopathy”, “Adrenergic signaling in cardiomyocytes” and “ECM-receptor interaction” pathways are unique for *OSF*.

We observe that 4 out of the 5 unique pathways of *OSF* are related to cardiomyopathies. A potential connection between *OSF* and cardiomyopathies is missing from the bibliography. However, in their review, Jiang *et al.* highlight drugs from various categories such as steroids, enzymes, cardiovascular drugs, antioxidants, vitamins, and microelements that seem to ameliorate (but not cure) the fibrotic effects of *OSF* (Jiang and Hu, 2009). In particular, they present cardiovascular drugs that were used in other studies against *OSF* including (i) pentoxifylline (Rajendran, *et al.*, 2006), (ii) buflomedil hydrochloride (Lai, *et al.*, 1995) and (iii) nylidrin hydrochloride (Sharma, *et al.*, 1987) as well as (iv) tea pigment (Li and Tang, 1998) all of which show temporary symptomatic relief. We suggest that a combinatorial treatment including a cardiovascular agent targeting unique highlighted cardiac-related pathways of the *OSF* disease should be further pursued.

In Figure 6, we present a similarity network based on common pathway terms among the 9 fibrotic diseases. *IPF* does not have any exclusive pathway nodes at the top-*5*% of its PathWalks results. *IPF* shares the most pathways with *IgG4-related Disease* (*14*/*15*) and the fewest with *OSF* (*8*/*15*). More information regarding the highlighted unique and in-common pathways among the 9 fibrotic diseases is shown in Supplementary Table 3. In Table 4, we present all pathways that were returned at the top-*5*% of the PathWalks results for at least 1 experiment. We also delineate entries at the top-*5*% of pathways with highest betweenness centrality and degree scores as well as entries at the top-*5*% results of a random PathWalks execution (without the use of a gene map). Here, we highlight pathways that were produced by our algorithm exclusively due to their association with the genetic information maps in green color.

**Figure 6:**
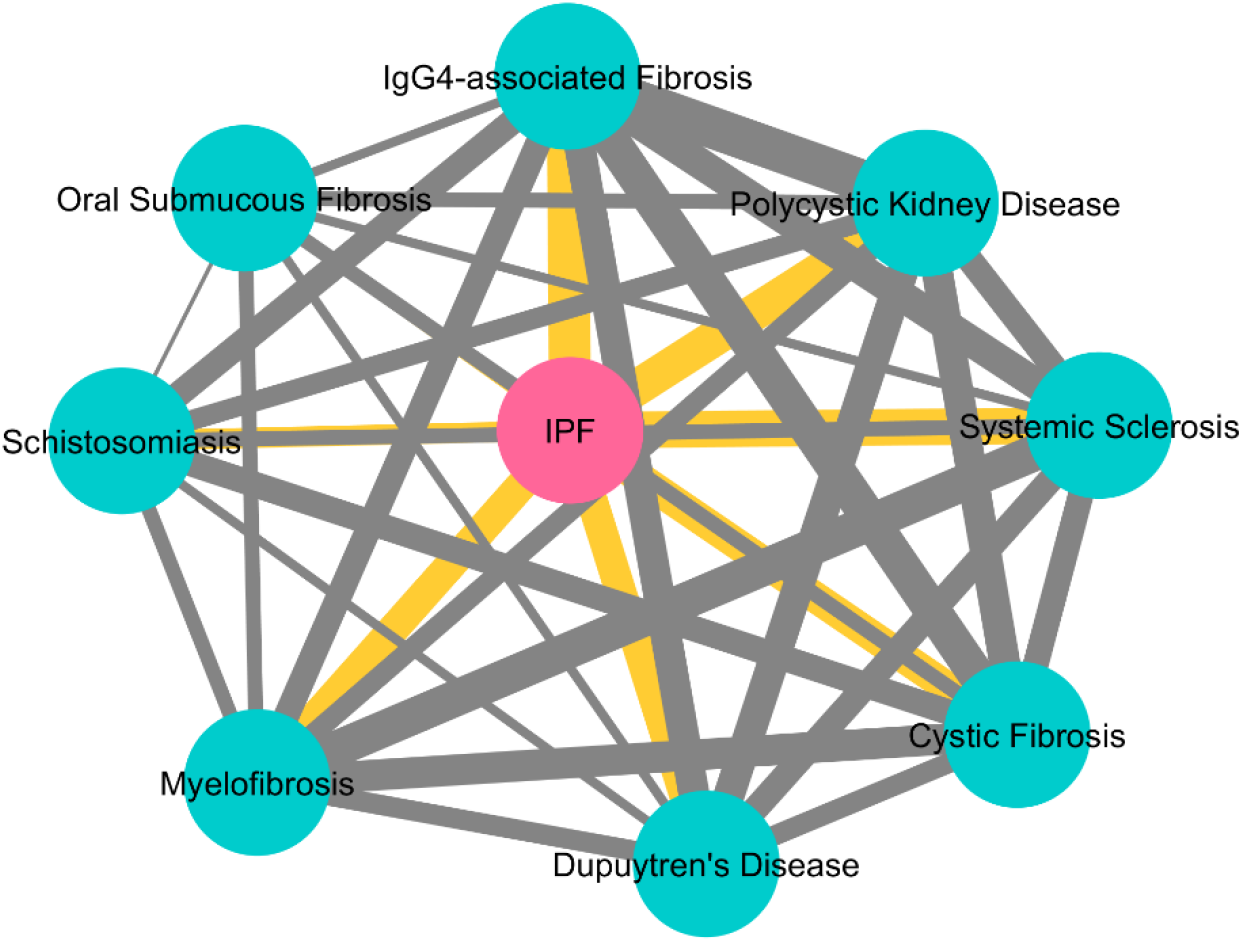
Similarity network among the 9 fibrotic diseases based on their common pathways as produced by the PathWalks algorithm. *IPF* does not have any unique pathway nodes at the top-*5*% of its PathWalks results. *IPF* has the most common pathways with *IgG4-related Disease* (*14*/15) and the least with *OSF* (*8/15*). The edge thickness is relative to the number of common pathways between any 2 diseases.

**Table 4:** Participation table of pathways at the top-*5*% of betweenness centrality, degree and random PathWalks (without gene map) result nodes. The 1st column of the table contains all pathways that reached the top-*5*% in at least 1 PathWalks experiment. We provide a check mark for each pathway at the top-*5*% of betweenness centrality, degree and random PathWalks lists respectively. We paint pathways that are completely highlighted by our algorithm due to their relation with each genetic information map in green.

| Top 5% Pathways' Table |  |  |  |
| --- | --- | --- | --- |
| Aggregated List of Top Pathways | Betweenness Centrality | Degree | Random PathWalks |
| Adrenergic signaling in cardiomyocytes |  |  |  |
| Alzheimer disease |  |  | ✓ |
| Antigen processing and presentation |  |  |  |
| Apoptosis | ✓ | ✓ | ✓ |
| Calcium signaling pathway | ✓ | ✓ | ✓ |
| cAMP signaling pathway | ✓ |  | ✓ |
| Cardiac muscle contraction |  |  |  |
| Cell adhesion molecules |  |  |  |
| Complement and coagulation cascades |  |  |  |
| Cytokine-cytokine receptor interaction |  |  |  |
| Dilated cardiomyopathy |  |  |  |
| ECM-receptor interaction |  |  |  |
| Fat digestion and absorption |  |  |  |
| Focal adhesion |  |  | ✓ |
| Glycerolipid metabolism |  |  |  |
| Herpes simplex virus 1 infection |  |  |  |
| Hypertrophic cardiomyopathy |  |  |  |
| IL-17 signaling pathway |  |  |  |
| Influenza A | ✓ |  | ✓ |
| MAPK signaling pathway | ✓ | ✓ | ✓ |
| Metabolic pathways | ✓ | ✓ | ✓ |
| Non-alcoholic fatty liver disease |  |  | ✓ |
| Oxidative phosphorylation | ✓ |  | ✓ |
| Pancreatic secretion |  |  |  |
| Pathways in cancer |  |  | ✓ |
| Pertussis |  |  |  |
| PI3K-Akt signaling pathway | ✓ | ✓ | ✓ |
| PPAR signaling pathway |  |  |  |
| Protein digestion and absorption |  |  |  |
| Regulation of actin cytoskeleton |  |  |  |
| Ribosome biogenesis in eukaryotes |  |  |  |
| RNA transport |  |  |  |
| Thermogenesis |  |  | ✓ |
| Toll-like receptor signaling pathway |  | ✓ | ✓ |

### Investigating Inhibiting Substances

Based on the procedure outlined in the Drug Repurposing segment of our Materials and Methods section, we collect repurposed drug candidates for the 9 respective fibrotic diseases through the CMap and L1000CDS^2^ DR tools. We then use our CoDReS re-ranking tool to extract the highest-scoring, structural-representative and inhibiting substances among the repurposed drugs of each disease. For *IgG4-related Disease,* we arbitrarily choose the top-*5* ranked drugs from each DR tool since there is no entry for this disease in CoDReS. In Table 5, we present the re-ranked candidate drugs for each disease.

**Table 5:** The re-ranked repurposed drugs for the respective fibrotic diseases. The 1^st^ column contains the disease names while the 2^nd^ column contains the names of the respective maladies in the database of CoDReS. The 3^rd^ column indicates the highest-ranked structural representative inhibitors returned by CoDReS and the 4^th^ column indicates their number. *IgG4-related Disease* is an exception since there is no matching entry in CoDReS. For this use case we present the top-*5* ranked drugs from each DR tool (CMap and L1000CDS^2^).

| Disease Name | CoDReS Disease Name | Top-Ranked Structural Representatives | #Top Ranked Drugs |
| --- | --- | --- | --- |
| *IPF* | *Idiopathic Pulmonary Fibrosis* | bi-2536, boldine, brazilin, brompheniramine maleate, budesonide, buparlisib, butein, celecoxib, ciglitazone, cinchonine, disulfiram, entinostat, epinephrine, hydrocortisone, icosapent, ipriflavone, memantine, methimazole, mk-2206, pd-0325901, propofol, propranolol, rizatriptan, scriptaid, sorafenib, sunitinib, testosterone, topotecan | 28 |
| *CF* | *Cystic Fibrosis* | acepromazine, apigenin, damnacanthal, delcorine, didanosine, edaravone, entinostat, fasudil, gefitinib, hydrocortisone, mercaptopurine, nafcillin, navitoclax, nilotinib, orantinib, oxybutynin, propofol, quipazine, sb-202190, staurosporine, terbutaline, trametinib, wortmannin, y-27632, ziprasidone | 25 |
| *SSc* | *Systemic Scleroderma* | erlotinib, gsk-1059615, hydrocortisone, l-690330, orphenadrine, radicicol, sb-431542 | 7 |
| *Myelofibrosis* | *Myelofibrosis due to another disorder* | alaprocate, anisomycin, brefeldin a, dasatinib, emetine, flurandrenolide, fludroxycortide, fluticasone, l-690330, narciclasine, nortriptyline, pd-0325901, pf-477736, procyclidine, regorafenib, reversine, ruxolitinib, sulpiride, tranlycypromine, trifluoperazine, xanthohumol, zalcitabine | 22 |
| *OSF* | *Oral Submucous Fibrosis* | cefotiam, cephaeline, dephostatin, digoxin, enzastaurin, decafluorobutane, xanthohumol | 7 |
| *Dupuytren's Disease* | *Dupuytren's Disease* | bi-2536, dasatinib, dicycloverine, fostamatinib, ibuprofen, l-690330, memantine, pha-665752, radicicol, triamcinolone, xylazine | 11 |
| *Polycystic Kidney Disease* | *Polycystic Kidney Diseases* | bosutinib, ipsapirone, loreclezole, picotamide, selumetinib, serdemetan, tretinoin, wortmannin | 8 |
| *IgG4-related Disease* | - | BRD-K92317137, brefeldin-a, cerulenin, dexketoprofen, fenpiverinium, gestrinone, importazole, indirubin, parthenolide, rhodomyrtosin | 10 |
| *Schistosomiasis* | *Schistosomiasis* | 1-benzylimidazole, canertinib, captopril, chloroquine, dapsone, dexamethasone, genipin, ibuprofen, iodophenpropit, levocabastine, mdl-28170, memantine, nadolol, parthenolide, phenytoin, propantheline, resorcinol, scriptaid, sunitinib, trametinib, triamterene, vemurafenib | 22 |

There are 2 drugs suggested by CoDReS that are common among 3 of the 9 fibrotic diseases. Hydrocortisone is highlighted as an anti-fibrotic drug candidate from the re-ranking procedure for *IPF*, *CF* and *SSc*. Similarly, memantine is highlighted in the use cases of *IPF*, *Dupuytren’s Disease* and *Schistosomiasis*. Hydrocortisone is a corticosteroid and its mode of action against fibrotic diseases has been studied for more than half a century now. It is mostly used for the temporal treatment of *OSF*. More specifically, Desa *et al.* report relief of symptoms in *OSF* patients by injecting the submucosal fibrotic areas with specific dosages of hydrocortisone depending on the stage of the fibrosis (DeSa, 1957). In a more recent study, Singh *et al.* describe how they have been clinically using optimal doses of a combination of hydrocortisone acetate and hyaluronidase for the medical treatment of *OSF* during a period of 20 years (Singh, *et al.*, 2010).

Hydrocortisone’s effect has also been studied against *Pulmonary Fibrosis* (*PF*). Xu *et al.* show that hydrocortisone decreases overexpressed fibrotic factors, such as TGF-β1, pulmonary type I collagen and inducible nitric oxide synthetase to normal levels in a rat model (Xu, *et al.*, 2007). Regarding *CF*, the study of Tepper *et al.* suggests that the application of intravenous hydrocortisone to the standard treatment of hospitalized infants with *CF*, produces a greater or a more sustained improvement in their lung function (Tepper, et al., 1997). In addition (Zachariæ and Zachariae, 1957) demonstrate the effects of hydrocortisone on various fibrotic skin maladies. Specifically, regarding *Dupuytren’s Disease,* they observe that the institution of hydrocortisone acetate at a mild stage keeps the tissue contracture stationary. Regarding *IgG4-related Disease*, Tanabe *et al.* propose a combination of hydrocortisone and thyroxine which result in clinical and laboratorial improvements against the disease (Tanabe, *et al.*, 2006). Finally, Mitani *et al.* study the effects of hydrocortisone in a UV-irradiated mouse model. The UV irradiation causes the appearance of fibroblasts and the accumulation of collagen in the model’s lower dermis. In (Mitani, *et al.*, 1999), they establish that hydrocortisone prevents both the fibrosis of lower dermis and the accumulation of ECM components.

Memantine is a low-affinity voltage-dependent uncompetitive antagonist of the N-methyl-D-aspartate receptors found in nerve cells and is used for the treatment of patients with moderate to severe *Alzheimer’s Disease*. A direct connection of memantine and fibrotic diseases does not exist in bibliography. However, the histopathological analysis by Abbaszadeh *et al.* show a successful pre-treatment of a heart failure rat model with memantine, that attenuates myocyte necrosis and fibrosis (Abbaszadeh, *et al.*, 2018). Similarly, Li *et al.* demonstrate that memantine attenuates lung inflammation in a bleomycin-induced acute lung injury mouse model (Li, *et al.*, 2015). They propose that further studies should be carried out to identify memantine’s mode of action regarding both the prevention as well as the treatment of lung fibrosis.

Based on the common re-ranked drug candidates from our analysis we derive a disease similarity network in Figure 7. The edge thickness is relative to the number of common re-ranked drugs between any pair of diseases. We calculate the significance of a disease sharing re-ranked drug candidates with *IPF*. In this regard, we use R’s hypergeometric test function *dhyper(x, m, n, k)* where (i) *x* is the number of shared CoDReS re-ranked drug candidates between *IPF* and another fibrotic disease, (ii) *m* is the cardinality of the intersection between the *IPF* CoDReS drugs (*28*) and the total, unfiltered drug entries that are produced by CMap and L1000CDS^2^ across all the experiments of a disease, (iii) *n* is the number of all the unfiltered repurposed drugs for the disease, excluding the shared ones with *IPF,* and (iv) *k* is the number of the CoDReS re-ranked drugs for the disease. For example, *CF* shares *3* (*x*) CoDReS re-ranked drug candidates with *IPF*, where *26* (*m*) out of the *28* re-ranked *IPF* drugs exist in the unfiltered outputs from the two DR tools for *CF*, *2448* (*n*) exist in the *CF* unfiltered DR outputs and were not highlighted for *IPF* through CoDReS and *25* (*k*) is the number of CoDReS-highlighted *CF* drug candidates. Therefore, the fact that *CF* shares 3, potentially anti-fibrotic, drug candidates (entinostat, hydrocortisone, propofol) with *IPF* is statistically significant (*p-value ≃ .002*). Based on this test, sharing 2 or 3 re-ranked drug candidates with *IPF* is a statistically significant (*p-value < .05*) similarity while sharing only 1 drug may happen due to randomness. *IPF* shares memantine, scriptaid and sunitinib with *Schistosomiasis*, bi-2536 and memantine with *Dupuytren’s Disease*, hydrocortisone with *SSc* and pd-0325901 with *Myelofibrosis*. In Supplementary Table 4, we offer detailed information regarding the common and unique highlighted candidate drugs among the 9 fibrotic diseases of interest.

**Figure 7:**
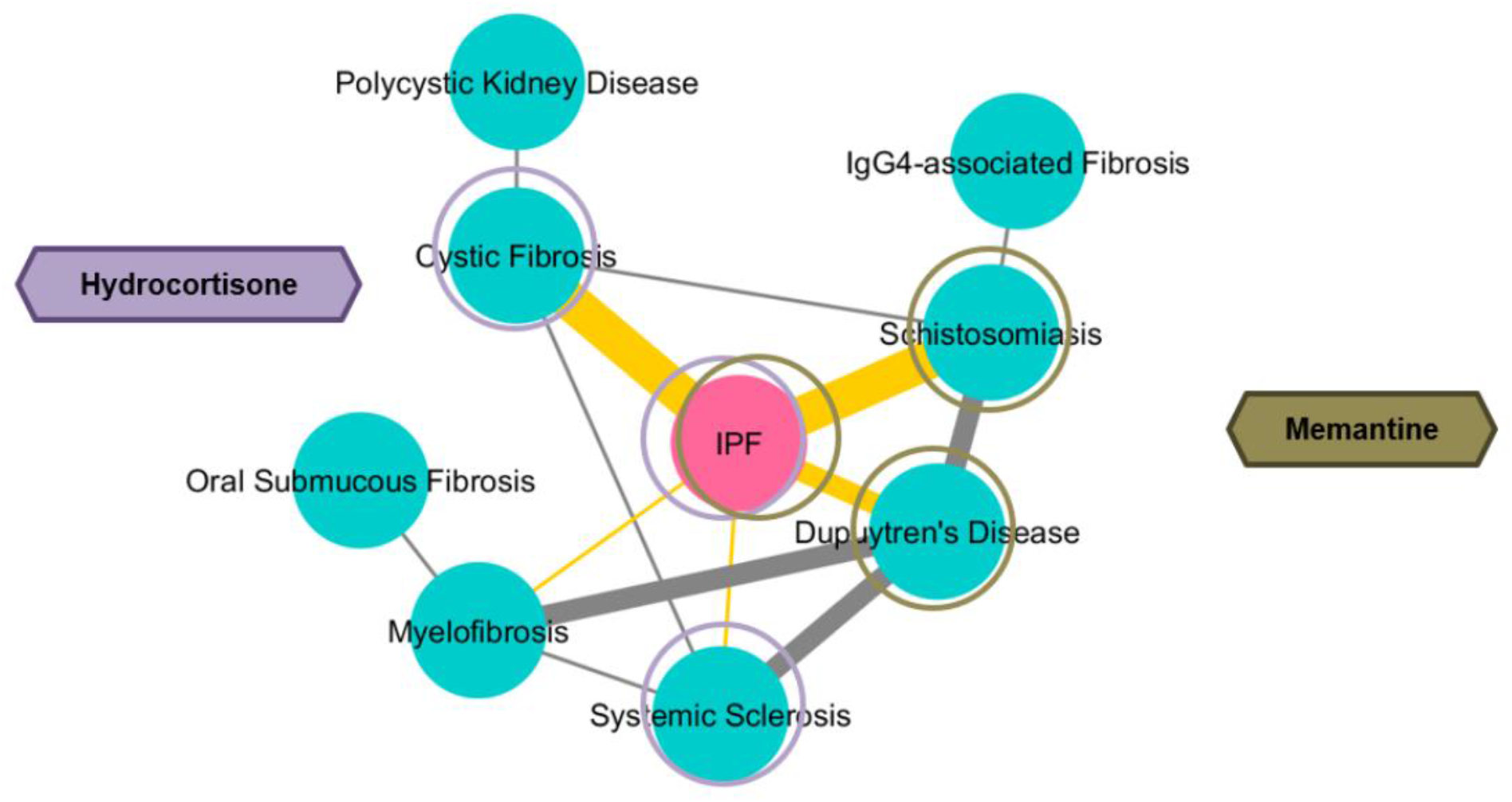
Disease-similarity network based on the common re-ranked drug candidates. The edge thickness is proportional to the number of common genes between pairs of diseases. *IPF* shares 3 drugs with *CF* and *Schistosomiasis*, 2 with *Dupuytren’s Disease* and 1 drug with *SSc* and *Myelofibrosis*. The fact that *IPF* shares 2 or 3 drugs with each of *CF*, *Schistosomiasis* and *Dupuytren’s Disease* is statistically significant based on a hypergeometric test. *SSc* and *Myelofibrosis* each share only 1 re-ranked drug with *IPF*, an event that might occur due to randomness. Hydrocortisone is highlighted by the re-ranking procedure for *IPF*, *CF* and *SSc* while memantine is highlighted in the use cases of *IPF*, *Dupuytren’s Disease* and *Schistosomiasis*.

In an effort to further screen the repurposed and re-ranked candidate drugs, we explore structural similarities among them and identify drugs that have previously failed in clinical trials against fibrotic diseases. The repoDB database contains entries with substances and their clinical trial status against the respective diseases. The 4 different drug status types of annotation in the database are: “Approved”, “Suspended”, “Terminated” and “Withdrawn”. We focus on drugs that have previously failed in clinical trials against any of the 9 fibrotic diseases of our study. We consider that drugs with “Suspended”, “Terminated” or “Withdrawn” indications have failed in the respective clinical trial. We place detailed information regarding drug names, respective fibrotic disease and clinical trial indication combinations, as found in repoDB, in Supplementary Table 5. We note that alemtuzumab (terminated entry against *SSc*) does not have a chemical structure file in PubChem (Kim, *et al.*, 2016), so we focus our analysis on the *19*-rest “failed” substances.

We use PubChem-available 2-dimensional structural data files (SDF) for the aforementioned *19* failed drugs and compile an aggregated SDF file. We then compile another SDF file containing the structures of all unique *121* CoDReS re-ranked drug candidates from this study. We use these 2 SDF files as input in the ChemBioServer “*Attach similar-only nodes to Network*” function. We use the SDF file of re-ranked drugs as a base network with an edge similarity threshold of *0.5* and the SDF file of failed drugs as a secondary list to infer structural similarity links towards the base network with an edge threshold of *0.7*. We choose the Tanimoto similarity metric since it is known to yield accurate results regarding cheminformatic similarity calculations (Bajusz, et al., 2015; Woodruff, et al., 1975). We then export the network created by ChemBioServer, remove drug nodes that are dissimilar to the failed drugs and visualize the remaining network in Cytoscape, as appears in Figure 8. With gray, we color the re-ranked drugs and with green, we color the failed substances. The edge thickness is commensurate to the structural similarity between any 2 compounds.

**Figure 8:**
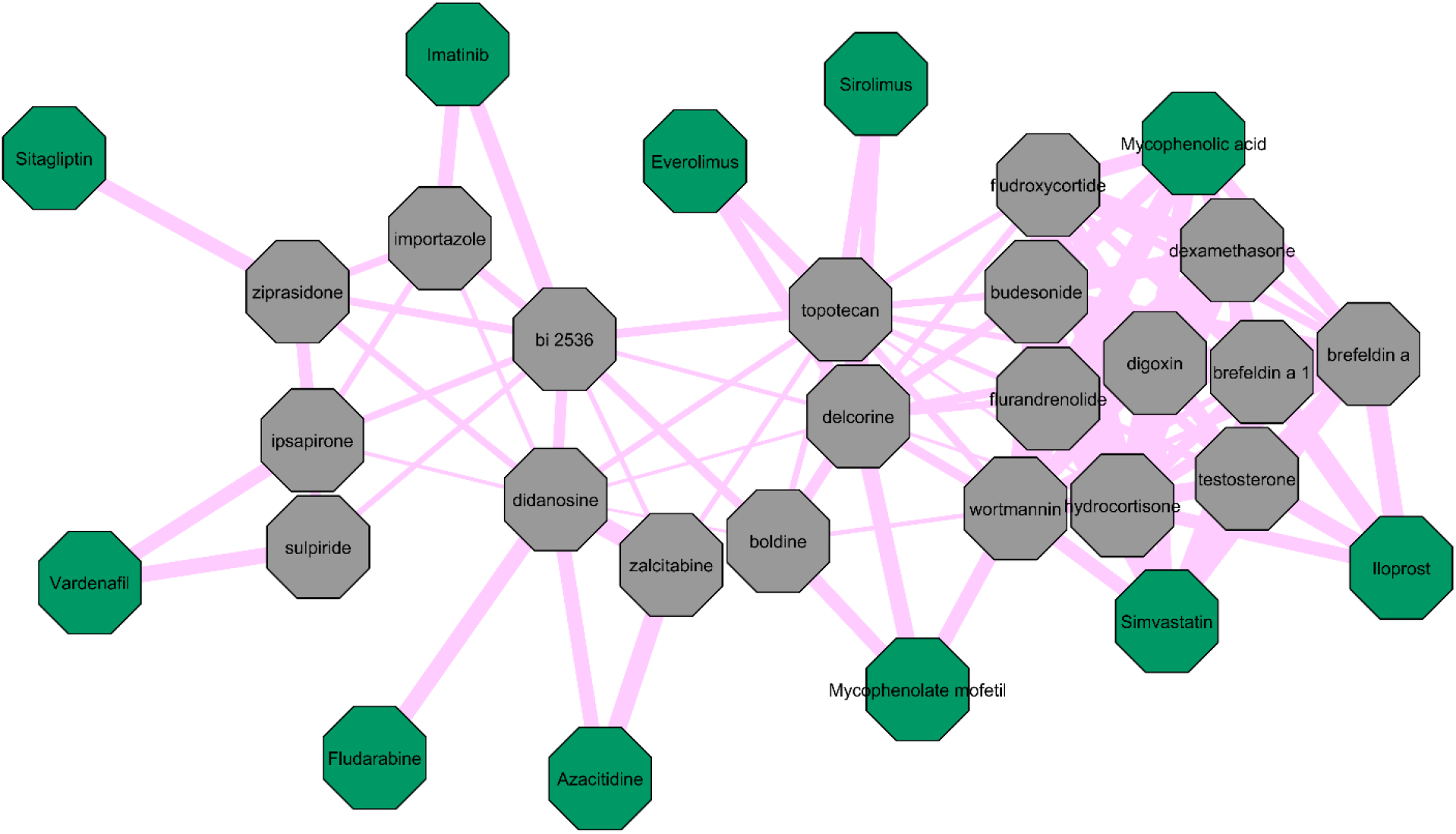
Structural similarity network of the CoDReS re-ranked drug candidates and drugs that have previously recorder as failed in clinical trials against fibrotic diseases. Re-ranked drugs that are dissimilar to the “failed” drugs have been removed from the network. We color the re-ranked drugs gray and the “failed” drugs green. The edge size is relative to the structural similarity between 2 compounds. Hydrocortisone, which has been re-ranked for 3 out of the 9 fibrotic diseases, has high structural similarity with iloprost, mycophenolic acid and simvastatin. We suggest giving lower testing priority to the depicted re-ranked substances due to their high similarity with already-failed drugs in clinical trials regarding fibrosis.

Figure 8 points out that hydrocortisone has high structural similarity with the “failed” drugs iloprost, mycophenolic acid and simvastatin. Hence, we suggest giving lower testing priority to hydrocortisone as well as to the rest of the substances of Figure 8 due to their high structural similarity with drugs that have already failed in clinical trials as far as fibrotic diseases are concerned. Nevertheless, we do not reject the possibility that even these potential drug choices may act as fibrotic inhibitors in conjunction with other drugs or at different doses and/or stages of fibrosis.

### Highlighting Anti-fibrotic Drug Candidates

Through our analysis pipeline, we settle for 121 total repurposed and re-ranked drug candidates with potential anti-fibrotic action. From those 121, 89 target at least 1 gene participating in at least 1 of the PathWalks highlighted pathways for the 9 fibrotic diseases. We focus on substances targeting *IPF* highlighted pathways that are directly associated with the gene map of *IPF*. There exist 3 such pathways: “Pancreatic secretion”, “Protein digestion and absorption” and “Complement and coagulation cascades” colored green in Figure 9. This figure zooms into the communities of the top-*5*% PathWalks-highlighted pathways for *IPF* and their functional connections. The edge weights are proportional to the number of times they were traversed by our PathWalks algorithm. The “Pancreatic secretion” and “Protein digestion and absorption” pathways are highlighted for *IPF*, *Dupuytren’s Disease*, *Polycystic Kidney Disease* and *IgG4-related Disease*. The “Pancreatic Secretion” pathway is also highlighted for *SSc*. The “Complement and coagulation cascades” pathway is highlighted in the cases of *IPF*, *CF*, *IgG4-related Disease*, *Polycystic Kidney Disease* and *Schistosomiasis*.

**Figure 9:**
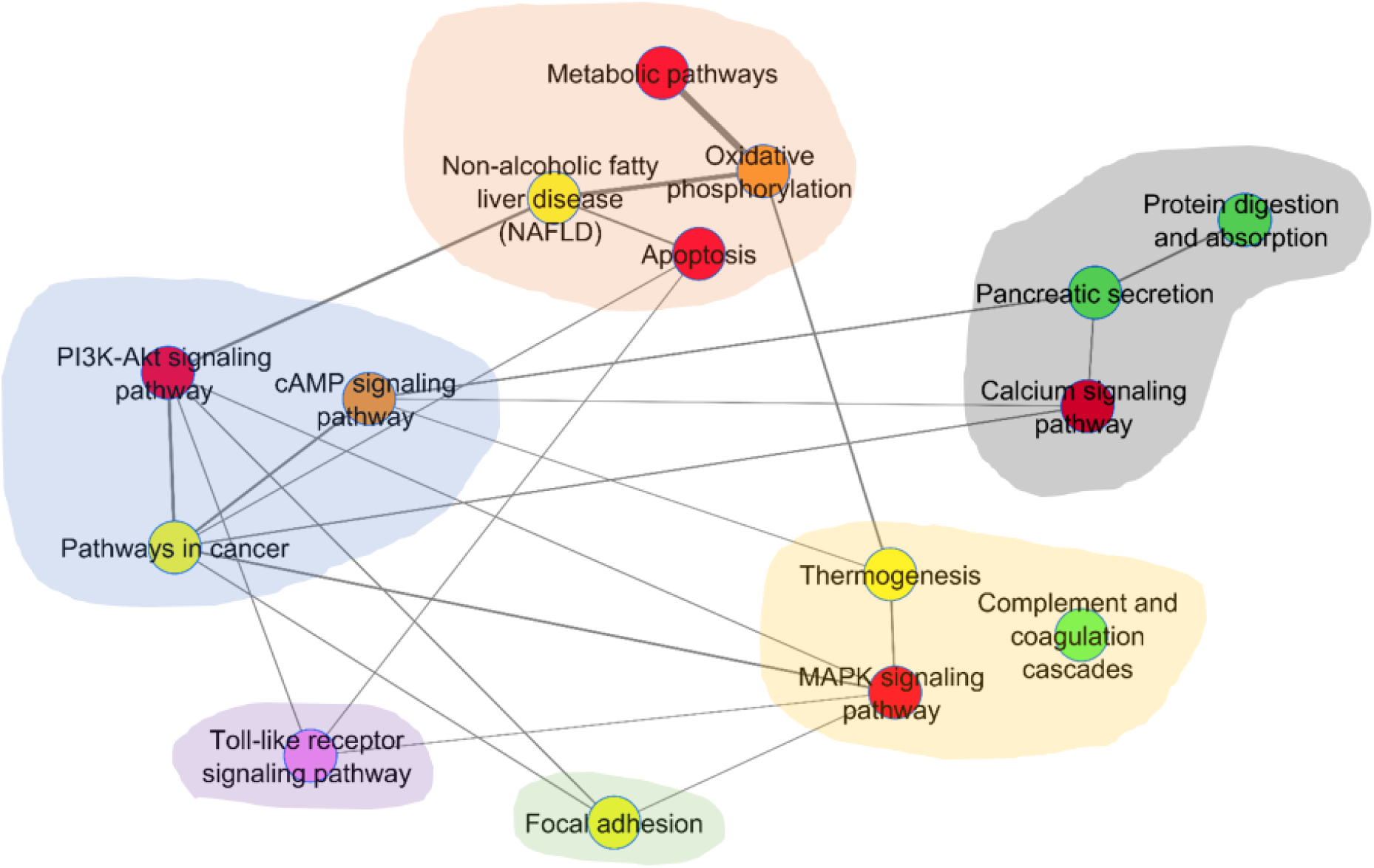
Pathway communities at the top-5% results for *IPF* as produced by PathWalks. “Pancreatic secretion”, “Protein digestion and absorption” and “Complement and coagulation cascades” are the 3 pathways that are highlighted exclusively due to their direct association with the genetic information network of *IPF* and are colored in green. The edge width is commensurate to the number of times an edge was traversed during the *IPF* PathWalks execution. The “Pancreatic secretion” and “Protein digestion and absorption” pathways are highlighted in *IPF*, *Dupuytren’s Disease*, *Polycystic Kidney Disease* and *IgG4-related Disease*. The “Pancreatic Secretion” pathway is also highlighted in *SSc*. The “Complement and coagulation cascades” pathway is highlighted in the use cases of *IPF*, *CF*, *IgG4-related Disease, Polycystic Kidney Disease and Schistosomiasis*.

There are 34 out of the 121 total re-ranked drugs that target at least 1 gene participating in the 3 “green” pathways of *IPF* in Figure 9. In Supplementary Table 6, we present all the combinations of re-ranked drugs, gene targets and biological pathways, where a drug has at least 1 gene target in the highlighted ‘green’ pathways. We bibliographically explore the 7 candidates that target at least 2 out of the 3 ‘green’ highlighted pathways of Figure 9 for potential anti-fibrotic mode of action, namely celecoxib, digoxin, captopril, ibuprofen, staurosporine, nafcillin, wortmannin.

Celecoxib is the only candidate targeting all 3 “green” pathways of *IPF*. Specifically, celecoxib is an anti-inflammatory drug that we identify in the CoDReS results for *IPF*. Celecoxib targets the *CA2* and *COL18A1* genes in “Pancreatic secretion” and “Protein digestion and absorption” pathways respectively and *PLAU* in “Complement and coagulation cascades”. We identify *PLAU* being over-expressed in the *Dupuytren’s Disease* experiments. The anti-fibrotic effects of celecoxib have been widely studied, especially in rat models and mainly regarding liver fibrosis. A number of such studies (Chávez, *et al.*, 2010; Ftahy, *et al.*, 2013; Gao, *et al.*, 2013; Paik, *et al.*, 2009; Wen, *et al.*, 2014) suggest the anti-fibrotic potential of celecoxib against hepatic fibrosis. On the other hand, contradictory studies (Hui, *et al.*, 2006; Liu, *et al*., 2009) demonstrate pro-fibrotic effects of celecoxib and their results show significant exacerbated liver fibrosis and hepatocellular necrosis in celecoxib-treated rats. However, Yu *et al.* show that celecoxib protects against liver inflammation in early stages of fibrosis even though it has no effect on the actual fibrosis (Yu, *et al.*, 2009). Similarly, Harris *et al.* observe that celecoxib treatment has no effect in the liver fibrosis of mice (Harris, *et al.*, 2018). In a cardiac hypertrophy study, Zhang and colleagues demonstrate that celecoxib treatment prevents collagen accumulation and suppresses *CTGF* expression in rats (Zhang, *et al.*, 2016). Collagen accumulation and overexpression of *CTGF* are established pro-fibrotic mechanisms. Nevertheless, and mostly due to contradicting studies, we suggest that other potential drug choices from our study should be prioritized instead of celecoxib for further experimentation against fibrosis.

Digoxin has been adopted for the treatment of atrial fibrillation and congestive heart failure (Hallberg, *et al.*, 2007). We identify digoxin as a repurposed drug in the use case of *OSF*. As we mention in the pathway community analysis paragraph, there are 4 unique cardiomyopathy-related pathways involved in *OSF*, a finding that needs further experimental validation. Digoxin targets the “Pancreatic secretion” and “Protein digestion and absorption” through 8 genes: *ATP1A1, ATP1A2, ATP1A3, ATP1A4, ATP1B1, ATP1B2, ATP1B3* and *FXYD2*. *ATP1A2* is over-expressed in *OSF* and under-expressed in *Dupuytren’s Disease* while ATP1B1 is over-expressed in *Dupuytren’s* and FXYD2 over-expressed in *Polycystic Kidney Disease*. Peckham and colleagues study the effects of digoxin on nasal potential difference in *CF* subjects with perturbed airway epithelium. Their results indicate that digoxin is unlikely to treat *CF* (Peckham, *et al.*, 1995). On the other hand, Moss *et al.* study the effects of digoxin in *CF* patients with heart failure complications. Their results show promise in cardiac process reversion even though it is not certain which treatment among respiratory care, vigorous antibiotic therapy or digoxin, attributed most to the survival (Moss, *et al.*, 1975). Contrary to the previous reports, the study of Coates *et al.* concludes that digoxin neither increases exercise capacity nor improves exercising cardiac function in *CF* patients with moderate to severe airway obstruction (Coates, *et al.*, 1982).

Disregarding *CF*, Haller *et al.* observe remarkable clearing of myocardial fibrosis by an affinity-purified digoxin antibody in rats with *Chronic Renal Failure* (Haller, *et al.*, 2012). We observe that digoxin has high structural similarity with the “failed” drugs simvastatin and mycophenolic acid as well as hydrocortisone, as Figure 8 indicates. Nevertheless, we suggest that digoxin should be further tested specifically against *OSF* due to the results of our study hinting towards a link between *OSF* and heart diseases.

Captopril is used for the treatment of hypertension and some types of congestive heart failure (Vidt, et al., 1982) and is highlighted from the re-ranking process of our analysis in *Schistosomiasis*. Captopril targets the *F2* and *SERPINE1* genes in the “Complement and coagulation cascades” pathway and the DPP4 gene in the “Protein digestion and absorption” pathway. We observe SERPINE1 being over-expressed in *Myelofibrosis* and *SSc* but under-expressed in *IPF*. There are various studies highlighting captopril’s anti-fibrotic mode of action in experimental rat models. Regarding lung fibrosis, Ghazi-Khansari’s histopathological findings show that captopril helps in a paraquat-induced lung fibrosis rat model (Ghazi-Khansari, et al., 2007) while Baybutt’s study shows that captopril limits fibrosis and interstitial pneumonia in a monocrotaline-induced lung fibrosis rat model (Baybutt, et al., 2007). Jalil and colleagues’ study shows attenuation of both interstitial and perivascular fibrosis of myocardium in a renovascular hypertension rat model (Jalil, et al., 1991). Notable reduction of fibrosis has also been observed in rat studies regarding kidney (Cohen, et al., 1996), liver (Ramos, et al., 1994) and colon fibrosis (Wengrower, et al., 2004). We should note that there are fewer studies with contradictory results about captopril’s action against fibrosis. Tuncer and others do not observe any inhibitory effects of captopril in a rat model with liver fibrosis (Tuncer, et al., 2003) while Okada’s team note that captopril significantly enhances myocardial fibrosis in an isoprenaline-induced ventricular fibrosis rat model (Okada, et al., 2010). However, due to the various studies supporting captopril’s anti-fibrotic action as well as the indications from our study, we suggest that captopril should be considered for further examination against fibrotic diseases and especially *IPF* due to a number of existing reports for its capacity to improve lung fibrosis.

Ibuprofen is an anti-inflammatory agent that is highlighted in the use cases of *Dupuytren’s Disease* and *Schistosomiasis*. Ibuprofen targets the *PLAT* and *THBD* genes in the “Complement and coagulation cascades” pathway and *CFTR* in the “Pancreatic secretion” pathway. PLAT is over-expressed in *SSc*. Carlile *et al.* demonstrate that ibuprofen is a *CF* transmembrane conductance regulator (CFTR) protein corrector and suggest that ibuprofen may be suitable in a *CF* combination therapy (Carlile, *et al.*, 2015). Additionally, Lindstrom *et al.* suggest that high doses of ibuprofen administered based on weight can treat polyposis in children with *CF* (Lindstrom, *et al*., 2007). There are 3 more studies (Konstan, *et al.*, 1995; Konstan, *et al.*, 2007; Lands, *et al*., 2007) in agreement that high doses of ibuprofen slow the progression of lung disease in *CF* without serious adverse effects. Therefore, ibuprofen should be further experimentally pursued in drug combinations against fibrosis.

Staurosporine targets the *PLA2G1B* gene in the “Pancreatic secretion” pathway as well as COL1A2 and *SLC1A1* genes in the “Protein digestion and absorption” pathway. In our analysis, we identify *SLC1A1* as being over-expressed in *Polycystic Kidney Disease*. *COL1A2* is under-expressed in *Polycystic Kidney Disease* but over-expressed in *IPF*, *IgG4-related* and *Dupuytren’s Disease*. We note that the *PLA2G1B* gene is under-expressed in 3 out of the 6 total *IPF* experiments and over-expressed in 1 of them. Staurosporine is returned as a repurposed drug for *CF*. However, due to its non-specific selectivity on kinase-binding, staurosporine has a high chance of undesirable side effects (Karaman, *et al.*, 2008; Tanramluk, *et al.*, 2009). Moreover, the study of Lindroos *et al.* shows that staurosporine upregulates *PDGFR-α* gene expression and its protein levels in pulmonary myofibroblasts of rats (Lindroos, *et al.*, 2001). The overexpression of *PDGFR-α* is known to induce myofibroblast hyperplasia during pulmonary fibrosis. Therefore, we consider that staurosporine is not a viable anti-fibrotic candidate.

We designate nafcillin, an antibiotic drug, as a repurposed and re-ranked candidate in *CF*. Nafcillin targets the *F2RL3* gene in the “Complement and coagulation cascades” pathway and the *PGA5* gene in “Protein digestion and absorption”. *F2RL3* is under-expressed in *IPF* experiments. We suggest that nafcillin’s mode of action against fibrosis should be explored via cell line and model experiments since there are currently no relevant studies.

Finally, wortmannin is a phosphoinositide 3-kinase inhibitor (Powis, *et al.*, 1994) and is highlighted by CoDReS for both *CF* and *Polycystic Kidney Disease*. Wortmannin targets the *F2R* and *SLC1A1* genes in the “Complement and coagulation cascades” and “Protein digestion and absorption” pathways respectively. There are a few studies regarding *SSc* that hint at wortmannin’s potential anti-fibrotic mode of action. Shi-Wen *et al.* report that wortmannin greatly reduces *SSc* patients’ fibroblast ability to contract a collagen gel matrix (Shi-Wen, *et al.*, 2004). In a later study, they add that wortmannin blocks the ability of rac1 protein to increase the expression of other profibrotic proteins (Shi-Wen, *et al.*, 2009). Parapuram and colleagues also highlight wortmannin’s capacity to reduce the expression of collagen type I, *SMA* and *CCN2* in dermal fibroblasts from *PTEN* knockout mice (Parapuram, *et al.*, 2011). The study of Zhi *et al.* describes that wortmannin significantly decreases the expression of collagen type I in adult rat cardiac fibroblasts (Zhi, *et al.*, 2013). On the other hand, Wang *et al.* show that wortmannin cancels the effects of cannabinoid receptor type 2 agonist including improving cardiac function, decreasing fibrosis and reducing inflammatory cytokine secretion and oxidative stress in a mouse model with infarcted heart (Wang, *et al.*, 2014). Lastly, Zhang and others also identify that wortmannin abolishes zinc’s anti-fibrotic capacity in a rat kidney tubular epithelial cell line (Zhang, *et al.*, 2016). In Figure 8, we observe that wortmannin has high structural similarity with the “failed” drugs simvastatin, mycophenolic acid and mycophenolate mofetil, as well as with the already discussed repurposed drugs of our study hydrocortisone and digoxin. Due to the contradictory results from the bibliography and wortmannin’s structural similarity with already “failed” drugs against fibrosis, we do not suggest prioritizing wortmannin in future anti-fibrotic experiments.

## Discussion

In this study, we identify genes, biological pathways and potential inhibiting drugs that are associated with fibrotic diseases. Through our PathWalks methodology we identify 7 common biological pathways that are associated with all 9 fibrotic diseases investigated in this paper. These include “Metabolic”, “Cancer”, “MAPK signaling”, “PI3K-Akt signaling”, “Non-alcoholic fatty liver disease”, “Oxidative phosphorylation” and “Calcium signaling” pathways. We do not identify any exclusive fibrotic mechanisms for *IPF* but each of the *Myelofibrosis*, *CF*, *SSc*, *Dupuytren’s Disease*, *Schistosomiasis* and *OSF* diseases have unique pathway entries that should be further explored as to their contribution in the pathogenesis of these diseases. We observe that the PathWalks results hint evidence for a potential common mechanism between *OSF* and cardiomyopathies. Therefore, we suggest that combinatorial regimens against *OSF* including cardiovascular agents should be looked at.

We use 2 DR tools (CMap and L1000CDS^2^) and the drug re-ranking tool (CoDReS) to identify and propose drug candidates with potential anti-fibrotic mode of action. We observe that hydrocortisone and memantine are returned in 3 out of the 9 fibrotic diseases of this study. Memantine’s action has not been studied against fibrotic diseases but, as mentioned in our respective Results section, there are hints for its potentially beneficial anti-fibrotic action (Abbaszadeh, *et al.*, 2018; Li, *et al.*, 2015). As far as hydrocortisone is concerned, low doses have shown to attenuate fibrosis especially in the early stages (DeSa, 1957; Xu, et al., 2007). However, we note that hydrocortisone has high structural similarity with drugs that have previously failed in clinical trials against fibrosis. Even though we suggest prioritizing drugs that are dissimilar to “failed” ones, we do not reject the possibility that even these candidates may be effective in combination with other drugs or at different dosages and/or stages of fibrosis.

In our effort, we have also examined the repurposed drugs of this study that appear to be the most promising as anti-fibrotic choices. We select substances that target at least 2 out of the 3 PathWalks highlighted biological pathways of *IPF* and are directly associated with the respective gene map. We explore bibliographic evidence for pro- or anti-fibrotic mode of action regarding these drug candidates. Combining the results of our analysis and the bibliography, we suggest that captopril and ibuprofen should be prioritized for future anti-fibrotic experiments. We also suggest that nafcillin, which hasn’t yet been studied against fibrosis, should be considered for *in vitro* and *in vivo* studies. Finally, digoxin should also be further explored, specifically regarding *OSF* since, as stated in the Results section, it appears to clear myocardial fibrosis and *OSF* shares multiple biological pathways related to myocardial diseases.

We observe the *LCN2* gene being differentially over-expressed in 4 out of the 9 studied fibrotic diseases. Bibliographic evidence shows correlation between *LCN2* and fibrosis in *SSc*, *Chronic Hepatitis C* and *Chronic Kidney Disease*. However, in other diseases such as NAFLD and cases of liver injury, *LCN2* cannot be correlated to the observed fibrosis. More experimentation is needed to better understand the role of LCN2 in fibrotic implications. We also identify FBLN1 being under-expressed in 4 out of the 9 fibrotic diseases and over-expressed in another. According to bibliography, even though *FBLN1* mRNA levels may decrease in *COPD* where small airway fibrosis occurs, on the protein level it is accumulated in the ECM. FBLN1 levels are increased in serum and bronchoalveolar lavage fluid of asthma patients (Lau, *et al.*, 2010) and in the plasma and lung tissue of *IPF* patients (Jaffar, *et al.*, 2014). Based on our results, we observe an association between *FBLN1* and fibrotic diseases; this is also corroborated in existing bibliography (Ge, et al., 2015; Hansen, et al., 2013; Jaffar, et al., 2012) regarding specific tissues (e.g., lung, myocardium). We suggest that further proteomics analyses should be performed to measure the quantity of the fibulin-1 translated protein in the related fibrotic tissues and its potential involvement in fibrosis.

Our work establishes gene, pathway and candidate drug similarities among *IPF* and the rest of fibrotic diseases. Specifically, *IPF* shares several terms with *Dupuytren’s Disease* having *35* common over-expressed and *16* common under-expressed genes, 2 common key pathways with direct association to the respective genetic information maps and 2 common identified drug candidates. *IPF* and *Myelofibrosis* share *28* under-expressed genes. *IPF* and *IgG4-related Disease* share *23* over-expressed genes and 3 key pathways. Finally, *IPF* shares 20 over-expressed genes and 1 key pathway with *SSc* and 3 drug candidates and 1 key pathway with *CF*. Our conjecture is that common treatments for *IPF* and the aforementioned diseases, especially *Dupuytren’s Disease*, should be pursued.

We provide all derived results for brevity in Supplementary Material. Supplementary Tables 1 and 2 present the top over- and under-expressed genes per disease. In Supplementary Table 3, we depict the top-*5*% ranked pathways from the PathWalks algorithm and in Supplementary Table 4, we show the CoDReS re-ranked drugs that we identify for each fibrotic disease. Supplementary Table 5 presents the indications for the “failed” drugs of repoDB regarding clinical trials against fibrotic diseases. Finally, Supplementary Table 6 links drugs to their gene targets and the respective highlighted pathways. We stipulate that results on the gene, pathway and drug levels from this study will be further pursued to gain a deeper understanding of the pathogenesis as well as potential regimens against fibrosis.

